# U-Net Model for Brain Extraction: Trained on Humans for Transfer to Non-human Primates

**DOI:** 10.1101/2020.11.17.385898

**Authors:** Xindi Wang, Xin-Hui Li, Jae Wook Cho, Brian E. Russ, Nanditha Rajamani, Alisa Omelchenko, Lei Ai, Annachiara Korchmaros, Stephen Sawiak, R. Austin Benn, Pamela Garcia-Saldivar, Zheng Wang, Ned H. Kalin, Charles E. Schroeder, R. Cameron Craddock, Andrew S. Fox, Alan C. Evans, Adam Messinger, Michael P. Milham, Ting Xu

## Abstract

Brain extraction (a.k.a. skull stripping) is a fundamental step in the neuroimaging pipeline as it can affect the accuracy of downstream preprocess such as image registration, tissue classification, etc. Most brain extraction tools have been designed for and applied to human data and are often challenged by non-human primates (NHP) data. Amongst recent attempts to improve performance on NHP data, deep learning models appear to outperform the traditional tools. However, given the minimal sample size of most NHP studies and notable variations in data quality, the deep learning models are very rarely applied to multi-site samples in NHP imaging. To overcome this challenge, we used a transfer-learning framework that leverages a large human imaging dataset to pretrain a convolutional neural network (i.e. U-Net Model), and then transferred this to NHP data using a small NHP training sample. The resulting transfer-learning model converged faster and achieved more accurate performance than a similar U-Net Model trained exclusively on NHP samples. We improved the generalizability of the model by upgrading the transfer-learned model using additional training datasets from multiple research sites in the Primate Data-Exchange (PRIME-DE) consortium. Our final model outperformed brain extraction routines from popular MRI packages (AFNI, FSL, and FreeSurfer) across a heterogeneous sample from multiple sites in the PRIME-DE with less computational cost (20s~10min). We also demonstrated the transfer-learning process enables the macaque model to be updated for use with scans from chimpanzees, marmosets, and other mammals (e.g. pig). Our model, code, and the skull-stripped mask repository of 136 macaque monkeys are publicly available for unrestricted use by the neuroimaging community at https://github.com/HumanBrainED/NHP-BrainExtraction.

## 1. Introduction

As the recent explosion of MRI data sharing in Nonhuman Primate (NHP) scales the amounts and diversity of data available for NHP imaging studies, researchers are having to overcome key challenges in preprocessing, which will otherwise slow the pace of progress (Autio et al. 2020; Milham et al. 2018; Messinger et al., 2021; Lepage et al. 2021). Among them is one of the fundamental preprocessing steps - brain extraction (also referred to as skull-stripping) (Seidlitz et al., 2018; Tasserie et al., 2020; Zhao et al., 2018). In both human and NHP MRI pipelines, brain extraction is often among the first early preprocessing steps (Esteban et al., 2019; Glasser et al., 2013; Seidlitz et al., 2018; Tasserie et al., 2020; Xu et al., 2015). By removing the non-brain tissue, brain extraction dramatically improves the accuracy of later steps, such as anatomy-based brain registration, pial surface reconstruction, and cross-modality coregistration (e.g., functional MRI, diffusion MRI) (Autio et al. 2020; Seidlitz et al. 2018; Acosta-Cabronero et al. 2008; Lepage et al. 2021). In humans, automated brain extraction tools have been developed (e.g., the Brain Extraction Tool [BET] in FSL, 3dSkullStrip in AFNI, the Hybrid Watershed Algorithm [HWA] in FreeSurfer, etc.) and easily inserted into a diversity of preprocessing pipelines (e.g. Human Connectome Project [HCP], fMRIPrep, Configurable Pipeline for the Analysis of Connectomes [C-PAC], Connectome Computational System [CCS], Data Processing & Analysis for Brain Imaging [DPABI]) (Cox, 1996; Craddock et al., 2013; Fischl, 2012; Glasser et al., 2013; Jenkinson et al., 2012; Ségonne et al., 2004; Xu et al., 2015; Yan et al., 2016). However, adaption for macaque brain extraction is significantly more challenging, as the data are often noisy due to the smaller brain and voxel sizes involved. The low signal-to-noise ratio (SNR) and strong inhomogeneity of image intensity compromise intensity-based brain extraction approaches, necessitating parameter customization to fit the macaque data (Messinger et al., 2021; Milham et al., 2018). For instance, a new option ‘-monkey’ has been developed to customize AFNI’s widely-used 3dSkullStrip function, which improves its performance for NHP data. Yet, the results are still mixed across datasets and often require further manual corrections (Fig 1).

**Figure 1.**
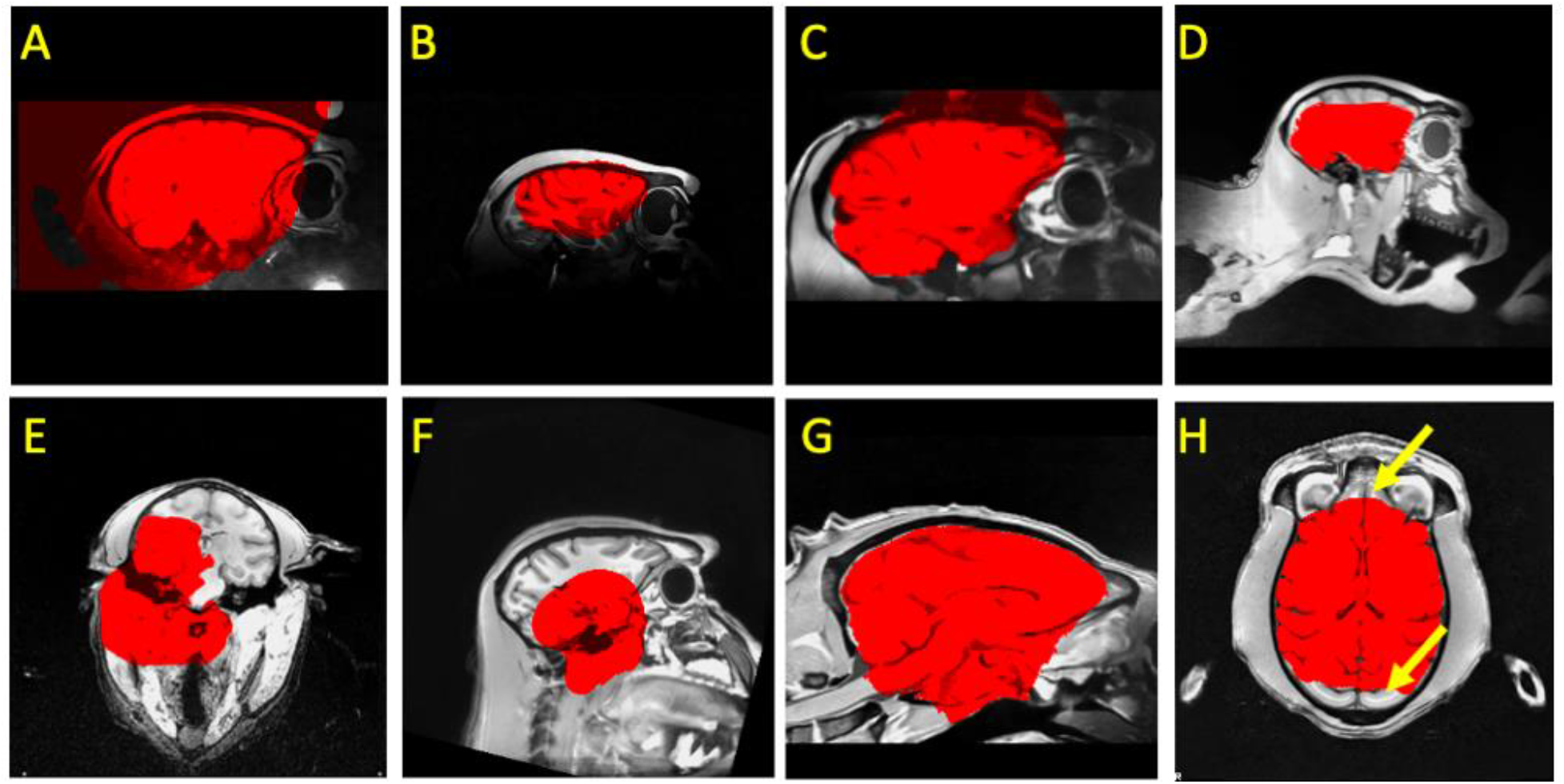
Examples of failures using traditional pipelines in NHPs. A) brain mask (red) extends beyond brain; B) a highly inhomogeneous T1w image with a mask that both misses parts of the brain and extends into the skull; C) an overly expansive mask for a NHP with a surgical implant (i.e. head holder); D) an incomplete mask for a brain that is not centered in the volume; E-H) other inaccurate masks.

In recent years, registration-based label transferring (i.e. template-driven) approaches have been proposed as a potential solution for NHP brain extraction (Jung et al., 2020; Lohmeier et al., 2019; Seidlitz et al., 2018; Tasserie et al., 2020). These approaches start by registering an individual’s anatomical image to the template in order to establish the deformation between the subject-specific head and template head. Once obtained, the transform is used to bring a template-based brain mask back to the individual space, where it can be used to extract the individual brain. The performance of such approaches heavily relies on the accuracy of the transformation and whether the template is representative of the individual data. As such, factors that can compromise the appropriateness or representativeness of the template for the specific individual dataset can decrease the utility of registration-based approaches. Examples where a template may not be representative include variations in macaque species (i.e. *M. mulatta, M. fascicularis*, etc.) and other NHPs species (e.g. macaque, marmoset, etc.), the field of view, age (e.g. infant, juvenile, aging), sex (e.g. thicker muscular tissue for male adult macaque), and surgical implants (e.g. with head-holder implants, or anatomical lesions). This issue might be further intensified for data from multiple study centers with different scan acquisitions and samples (Fig 1). In addition, non-linear registration (e.g. ANTs, 3dQwarp) for high-resolution images in brain extraction step is relatively time-consuming (e.g. over hours) and limits the computational efficiency of NHP pipelines.

Recognizing the continued challenges of brain extraction, researchers in human literature have begun to leverage deep learning models as a potential solution. In humans, a growing number of studies have demonstrated the ability of convolutional neural network (CNN) models for brain extraction, as well as tissue segmentation (Henschel et al., 2020; Lyksborg et al., 2015; Rehman et al., 2018; Snehashis Roy et al., 2018; Yogananda et al., 2019). Across studies, training and validation datasets in humans have included hundreds, and in some cases, even thousands of datasets, to ensure accurate performance and avoid overfitting. Once trained, the models have proven to be able to perform highly accurate extraction for new datasets in a matter of seconds. With rare exceptions, the NHP field does not possess datasets close to the multitudes used for training in humans. A recent study did, however, successfully implemented a CNN model (i.e. Bayesian SegNet) for brain extraction using a relatively smaller sample in macaques collected at a single site (N=50), suggesting that such training sets may not need to be as large as expected based upon these preliminary human studies (Zhao et al., 2018). However, the large majority of NHP studies use notably smaller sample sizes (i.e., 2-10). While combining data from multiple studies could be a solution, it is important to note the NHP literature tends to have substantially greater variability in imaging protocols than its human counterpart.

The present work attempts to overcome the challenges at hand for NHP imaging by developing a generalizable macaque brain extraction model that can handle data from previously untrained protocols/sites with high accuracy. To accomplish this, we leveraged a transfer learning U-NET strategy, which explicitly aims to train a model for one purpose, and then extended its utility to related problems. In the present case, we trained our model on a human sample (n = 197), and then treated a nonhuman sample (n = 2 for six sites) as the transfer dataset - a strategy that exploits the similarity of human and non-human brain structure. Upon successful demonstration of the ability to transfer between species, we then evaluated the transfer of the updated model to untrained sites in the PRIMatE Data Exchange. Finally, we improved the generalizability of our model by adding a single macaque sample from each of the additional 7 sites (for a total of N=19). We released our pre-trained model, code, and the brain masks outcomes via the PRIMatE Resource Exchange (PRIME-RE) consortium (Messinger et al., 2021; Milham et al., 2018).

## 2. Methods

### 2.1. Human Sample

We made use of an open available human brain extraction sample as an initial good-standard training dataset (Puccio et al., 2016). Data were collected as a part of the Enhanced Rockland Sample Neurofeedback Study (N=197, 77 female, age=21-45) (McDonald et al., 2017). Anatomical images data were acquired from a 3T Siemens Trio scanner using a 12 channel head matrix (T1-weighted 3D-MPRAGE sequence, FOV=256×256mm^2^, TR=2600ms, TE=3.02ms, TI=900ms, Flip angle=8°, 192 sagittal slices, resolution=1×1×1 mm^3^). T1w images were skull-stripped using a semi-automatic iterative procedure that involved skull-stripping all of the data using BEaST (brain extraction based on nonlocal segmentation technique) (Eskildsen et al., 2012) and manually correcting the worst results. Corrected brain masks were added into the BEaST library and the procedure was repeated until the process converged. The results of this procedure underwent an additional manual inspection and correction procedure to identify and fix any remaining errors.

### 2.2. Macaque Sample

The MRI macaque data used in the present study are publicly available from the recent NHP data-sharing consortium – the non-human PRIMate Data-Exchange (PRIME-DE) (Milham et al. 2018), which includes 136 macaque monkeys from 20 laboratories. We selected one anatomical T1w image per macaque in our analyses. The detailed description of the data acquisition of the magnetization-prepared rapid gradient echo (MPRAGE) image for each site was described in the prior study and PRIME-DE website (https://fcon_1000.projects.nitrc.org/indi/indiPRIME.html, Milham et al., 2018). As the sample size is relatively small in most of the sites (N≤6 for 13 out of 20 sites), we selected six sites which have no less than eight macaque monkeys collected as the first dataset pool for manual edits, model training, and testing (East China Normal University Chen [ecnu-chen], Institute of Neuroscience [ion], Newcastle University Medical School [newcastle], University of Oxford [oxford], Stem Cell and Brain Research Institute (sbri), and the University of California, Davis [ucdavis]). The details of the sample and MRI acquisition are specified in Milham et al ((Milham et al., 2018), Table 1). In total, eight macaque monkeys per site were selected to create the manually-edited ‘ground truth’ dataset to train and evaluate our model (**Macaque Dataset I**, N=48). To optimize the generalizability of our model to other sites, we also manually edited an additional seven macaques from seven sites (one per site: East China Normal University Kwok [ecnu-k], Lyon Neuroscience Research Center (lyon), Mount Sinai School of Medicine - Siemens scanner [mountsinai-S], National Institute of Mental Health - Messinger [nimh1], Netherlands Institute for Neuroscience [nin], and Rockefeller University [rockefeller]) to create **Macaque Dataset II** (N=7). The rest of the PRIME-DE macaques across all 20 sites were used as an additional hold-out testing dataset (N=81). Of note, our main model was built based on the MPRAGE data. We further made use of the magnetization-prepared two rapid acquisition gradient echoes (MP2RAGE) images from site-UWO-MP2RAGE (N=3) to extend our model to facilitate the brain extraction for MP2RAGE data. All animal procedures were conducted in compliance with the animal care and use policies of the institution where the data was collected.

### 2.3. Preprocessing

To improve the quality and homogeneity of input data across different sites, a minimal preprocessing was carried out for all anatomical images to remove the salt-and-pepper noise and correct the intensity bias. Specifically, we first re-conformed all T1w images into RPI orientation and applied a spatially adaptive non-local means filtering to remove the ‘salt-and-pepper’ noise (DenoiseImage in ANTs) (Buades et al., 2011). Next, we performed the bias field correction to normalize image intensities (N4BiasFieldCorrection in ANTs) (Tustison et al., 2010). The preprocessed images were served as inputs for all the brain extraction approaches.

### 2.4. Traditional Methods and Manually Edited Masks

To compare our deep learning models with state-of-the-art methods for brain extraction, we employed five widely-used skull stripping pipelines implemented in commonly used MRI packages (AFNI, ANTs, FSL, and FreeSurfer) (Avants et al., 2009; Cox, 1996; Fischl, 2012; Jenkinson et al., 2012). Specifically, we tested three intensity-based approaches (FSL BET, FreeSurfer HWA, and AFNI 3dSkullStrip) and two template-driven pipelines (Flirt+ANTS and AFNI @animal_warper) (Jung et al., 2020; Seidlitz et al., 2018; Tustison et al., 2020). The command and parameters of intensity-based approaches were selected based on the experiments and suggestions from the prior studies as follows (Xu et al., 2019; Zhao et al., 2018). 1) FSL ‘bet’ command with a smaller fractional intensity threshold (-f 0.3, denoted as “FSL”) and 2) with the vertical gradient in fractional intensity threshold and the head radius setting (-f 0.3 -g -0.5 -r 35, denoted as “FSL+”), 3) FreeSurfer ‘mri_watershed’ command with default settings and 4) using atlas information and brain radius setting (30 mm), 5) AFNI ‘3dSkullStrip’ command with NHP specific option ‘-monkey’ and shrink factor = 0.5. Template-driven approaches in both Flirt+ANTs and AFNI @animal_warper pipelines were performed by first applying a linear registration to transform the individual head to the template head, followed by nonlinear registration. Next, the template brain mask was transformed back into the individual space to obtain the individual brain mask. Specifically, the Flirt+ANTs pipeline uses ‘flirt’ and symmetric diffeomorphic image registration (SyN) (Avants et al., 2008) for linear and nonlinear registration. Of note, we used FSL ‘flirt’ rather than ANTS linear registration because ‘flirt’ is faster and performed better in our initial tests on NHP samples. AFNI @animal_warper uses 3dAllineate and 3dQwarp to compute affine and nonlinear alignments. The same NIMH Macaque Template (NMT) was used in the Flirt+ANTs and AFNI @animal_warper pipelines (Jung et al., 2020; Seidlitz et al., 2018). The details of the parameters and the approximate processing time are shown in Table S1. The Macaque Dataset I and II were manually edited by well-trained experts (J.W.C, A.K, and T.X.) using the best output from the traditional approaches as initial masks in ITK-SNAP (http://www.itksnap.org/pmwiki/pmwiki.php)(Yushkevich et al., 2006).

### 2.5. Train, Update & Evaluation Workflow for Deep Learning Models

#### 2.5.1. Overview

Figure 2 illustrates the overall analytic flow chart of the present study. First, we established a skull-stripping model using the human dataset, aiming to provide an initial pre-trained model to facilitate the transfer-learning from humans to macaques. The human NKI-RS dataset (N=197) was split into a training set (N=190) and a validation set (N=7). We used the training set to train the U-Net model for 10 epochs (i.e. the full training dataset passed through the complete neural network 10 times) and selected the best epoch as the human model based on the performance (i.e. dice coefficient) in the validating set. Next, we transferred the pre-trained human model to build the macaque model using Macaque Dataset I. Specifically, for each of six sites, we randomly selected 2 macaque monkeys as the training set, 1 macaque as the validation set and 5 macaques as the testing set. We also used all macaques across six sites in Macaque Dataset I to create the merged training (N=12), validating (N=6), and testing (N=30) sets.

**Figure 2.**
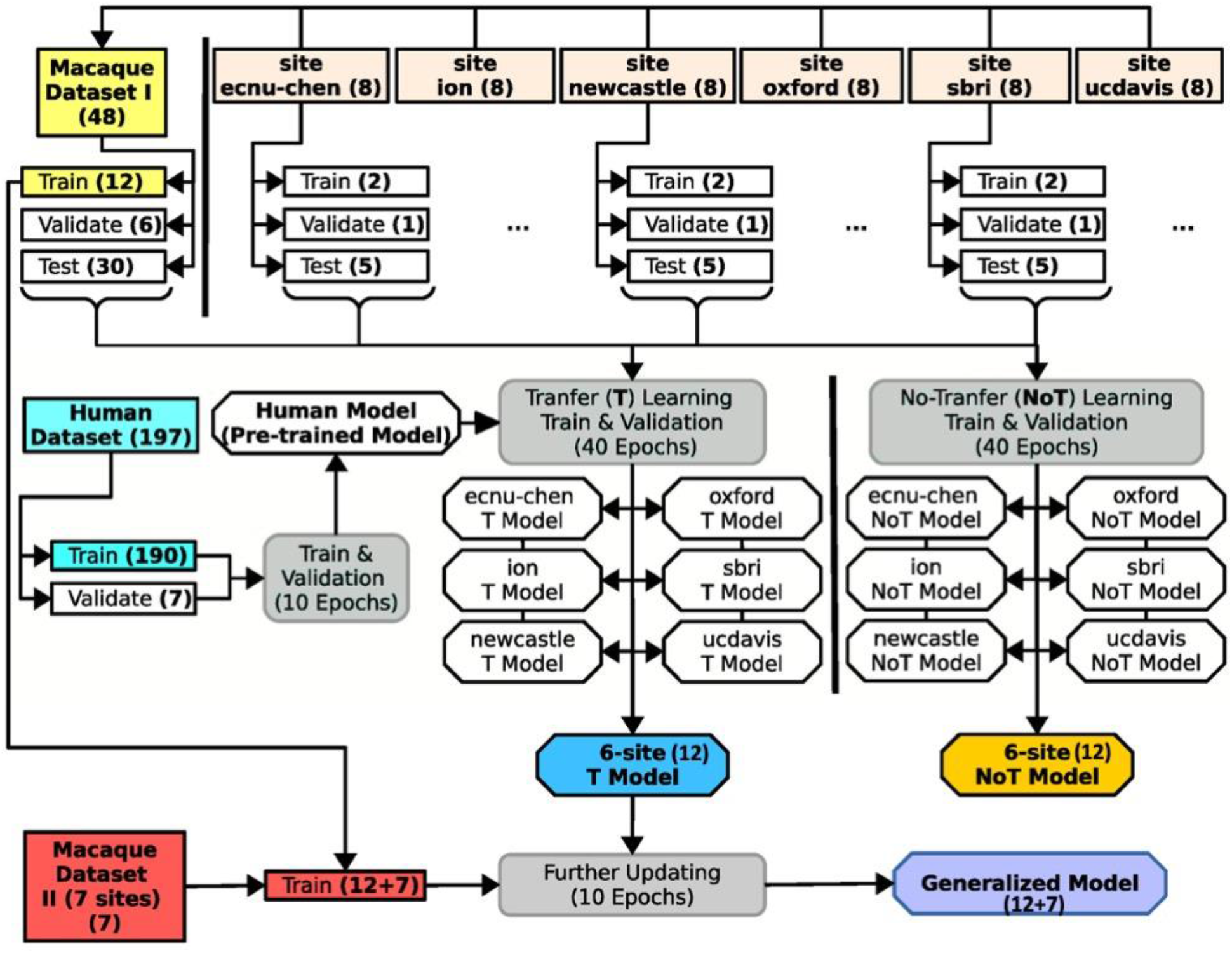
Schematic of the U-Net model’s training, transfer-learning, and validation steps. The datasets are highlighted in square boxes, models are in hexagonal boxes, and training and validating process (including transfer-learning) are in round boxes. The human dataset (light blue box, middle left) was used to pretrain the U-Net model for transfer learning. Macaque Dataset I (yellow box, upper left) included 12 subjects from 6 sites to train the model. Another 30 subjects from these sites were used to evaluate versions of this model with (T-model; blue angled box) and without (NoT-model; yellow angled box) transfer-learning from the human dataset. Macaque Dataset II (red box, lower left) was then added to the training set to generate a generalized model (purple angled box) for brain extraction on macaque data. Numbers in parentheses are subjects unless otherwise specified.

Transfer-learning from human to macaque was carried out for each site as well as for the merged data. We calculated 40 transfer-training epochs and selected the epoch that had the best performance in the validation set as the transfer-learning model for each site and the merged samples (refer to the U-Net T model). We also created macaque models that were trained only on the macaque data using the same training (N=12) and validation (N=6) data for each of the six sites and the merged samples (refer to the non-transfer-learning model, i.e. U-Net NoT model). We evaluated and compared the performance between the site-specific T model and the NoT model in testing sets. We also compared the U-Net T and NoT model to the traditional pipelines using the held-out test set from Macaque Dataset I. To improve the generalizability of the U-Net model to fit more macaque data from other sites, we further upgraded the U-Net transfer-learning model using both Macaque Dataset I (N=12) & II (N=7) to generate the final generalized model (referred to as generalized U-Net 12+7 model). To evaluate the model performance, we applied the U-Net transfer-learning model, generalized 12+7 model, and traditional pipelines to all the T1-weighted images from PRIME-DE (136 macaques). Expert ratings (details in the Model Evaluation section) were conducted to evaluate whether brain extraction was successful.

#### 2.5.2. Neural Network Model (U-Net)

We used a convolutional neural network (CNN) model, U-Net (Ronneberger et al., 2015) for brain extraction. The model was built using an open resource machine learning package (PyTorch: https://pytorch.org). Briefly, the preprocessed 3D T1w images were first resampled into slices along the axial, sagittal, and coronal planes. The U-Net model predicts the brain tissue in each slice and then merges all slices to obtain a 3D brain mask. Here, we focused on the architecture of the U-Net model (Fig. S1) and illustrated the details of how the U-Net model identifies the whole brain tissue in the training, validation, and testing processes in the next section.

As shown in Fig. S1, the U-Net model consists of a contraction (i.e. encoding) and an expansion (i.e. decoding) path; each includes five convolution units (Ronneberger et al., 2015). Of note, for a given slice resampled from the T1 images, the input of the U-Net model also included its neighboring slices (i.e. 3 slices in total) as an input (dimension: 3×256×256 blocks). Next, for each convolution unit, two 3×3 convolution layers are built and each is followed by a batch normalization and a leaky rectified linear (ReLU) operation (Fig. S1, blue arrow). In the encoding step, a 2×2 max pooling with stride 2 was adopted for down-sampling data from the upper unit to the lower unit. We used 16 feature channels in the initial unit, and doubled every unit. The expansive path consists of four up-convolution units. For each up-convolution unit, a 2×2 up-convolution and ReLU operation were applied to the lower unit to yield the feature map for the current unit. This feature map was then concatenated with the feature map at the same level in the contracting path to generate the combined feature map. Similar 3×3 convolution layers with batch normalization and ReLU operation were then performed on the feature maps at each up-convolution unit. Next, we employed a 1×1 convolution layer at the upper un-convolution unit to map the final feature maps to a two-classes map. Finally, a SoftMax layer was used to obtain the probability map for brain tissue. Of note, the initial weights of the convolution and up-convolution layers were randomly selected using a Gaussian distribution *N*(0, 0.2), and the initial bias of layers was set to 0. For the transfer-learning and model-upgrading model, the parameters from the pre-trained model were used in the initial setting.

#### 2.5.3. Model Training and Validating Procedure

In Figure S2, we illustrated the training procedure of how 3D T1w data was processed for the U-Net model. First, we normalized the intensity of the preprocessed T1w image in the range 0 to 1 across all voxels. We then resampled the T1w volume to a 3D intensity matrix where the highest sampled dimension of the T1w volume was forced to be rescaled to 256. In the example shown in Fig. S2, the initial T1w matrix of 176×176×96 was rescaled to a 256×256×140 matrix. Next, for each slice along the axial, sagittal, and coronal direction, we generated a 3-slice block; the slice and its two neighboring slices. As a result, we obtained 254, 254, and 138 blocks for axial, coronal, and sagittal directions respectively. Next, we conformed each of the 3-slice blocks into a 3×256×256 matrix. When the dimension of the slice plane was less than 256, we filled the matrix with zeros. In total, 646 conformed blocks were generated for the T1w image. Similarly, we processed the manually edited brain mask of the T1w image and generated 646 corresponding blocks. After that, we used the above U-Net model to estimate the probability of brain tissue for each T1w block and calculated the cross-entropy (PyTorch function: CrossEntropyLoss) between the probability map and the ‘ground truth’ map (Ketkar, 2017) as the model cost. A stochastic optimization (learning rate=0.0001, batch size=20) was then used for backpropagation (Kingma and Ba, 2014).

To evaluate and select the model from the training epochs, we used the probability map generated from the U-Net model to create the final predicted brain mask for each epoch, and then examined the Dice coefficient (see equation in section 2.6 below) between the predicted mask and the ‘ground truth’ mask (Fig. S3). Specifically, we processed the validation T1w images into 3-slices blocks following the above procedure. Next, we used the U-Net model at each training epoch to estimate the probability map for each block. All the probability maps (646 blocks) were then combined along the axial, sagittal, and coronal direction and yielded an averaged 3D probability matrix (256×256×256). After that, we rescaled and cropped the matrix back to the original voxel dimension to create a probability volume for the given T1w image. Finally, we thresholded (>0.5) this probability volume to obtain the predicted brain mask. The Dice coefficient between the predicted mask and the ‘ground truth’ mask was computed for each epoch during training. The epoch which showed the highest Dice coefficient was then selected as our final model.

### 2.6. Model Evaluation

We carried out a quantitative examination in the testing set of Macaque Dataset I and evaluated the degree to which methods provided more similar brain masks as compared to the manually edited ‘ground truth’. Specifically, we calculated Dice coefficients (Dice) (Sørensen, 1948) between the predicted mask and the manually edited ‘ground truth’ using the following equation:

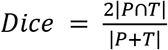

where |…| represents the total number of voxels in a mask and *P* and *T* are the predicted and ‘ground truth’ masks, respectively. Dice equals 1 when the predicted mask and ‘ground truth’ mask are perfectly overlapped. In addition, we also calculated the voxel-wise false-positive (FP), and false-negative (FN) rate to examine where the predicted mask falsely includes non-brain tissue (higher FP) or misses any brain tissue (higher FN). For each voxel in a given macaque, we tested whether the voxel is falsely assigned as brain tissue (FP=1) or falsely assigned as non-brain tissue (FN=1) in individual space. Next, we transferred the individual FP and FN map to the NMT space by applying affine and warp transforms. Linear affine here was created by aligning the manually skull-stripped brain to the NMT brain (‘flirt’) and nonlinear warp was generated by registering each individual’s head to the NMT head (‘fnirt’). The FP and FN maps were averaged across macaques for each brain extraction approach in the NMT space.

We also employed a qualitative evaluation to compare the success rate across different brain extraction approaches for PRIME-DE data without the ‘ground truth’. Three experts with rich experience in imaging quality control visually rated the brain masks (J.W.C., X.L., and T.X.). First, each expert independently reviewed all the images and rated them with four grades, i.e. ‘good’, ‘fair’ (the brain tissue was identified with a slightly inaccurate prediction at the edge), ‘poor’ (i.e. most of the brain regions are identified but with significant errors, in particular missing brain tissue at the edge), and ‘bad’. ‘Good’ and ‘fair’ rating were considered as a success while ‘poor’ and ‘bad’ scores were recognized as failures. The brain masks with inconsistent ratings (including good vs. fair, fair vs. poor, and poor vs. bad) among experts were reviewed and discussed for a final consensus rating.

## 3. Results

### 3.1. The Convergence of Human U-Net Model

To acquire a pre-trained model for macaque samples, we first trained a U-Net model using the human samples. Fig. S4 demonstrates the sum of loss on the training set and the mean Dice coefficients across human participants on the validation set for each epoch. After the first epoch, the loss decreased steeply and the mean Dice coefficient reached above 0.985. After that, the mean Dice coefficient gradually improved, showing its highest value (0.9916±0.0012) after the 9th epoch. To avoid over-fitting the human samples for the subsequent macaque training, we only carried out 10 epochs and selected the model after the 9th epoch as the pre-trained model for transfer-learning to the macaque samples.

### 3.2. Comparison Between Models with and Without Transfer-learning

We first evaluated the site-specific models with and without transfer-learning on Macaque Dataset I. For each site and for the merged sample across the six sites, the loss converged faster for the transfer-learning model than the model without transfer-learning (Fig. 3, left); the loss had nearly reached its minimum after 2 epochs. In addition, the Dice coefficients were more stable for transfer-learning models in the validation sets (Fig. 3, center). Of note, the mean Dice coefficients slowly dropped after 20 epochs in the validation set across three sites (i.e. ecnu-chen, ion, and sbri), which may reflect overfitting of the model selection on small training samples (Fig. 3, center). Nevertheless, the mean Dice coefficients still remained stable in the testing set after the first epoch (Fig. 3, right). In addition, compared to the model trained solely on macaque samples, the transfer-learning model also showed lower variation across testing macaques, suggesting its potential generalizability across macaque samples (Fig. 3, right).

**Figure 3.**
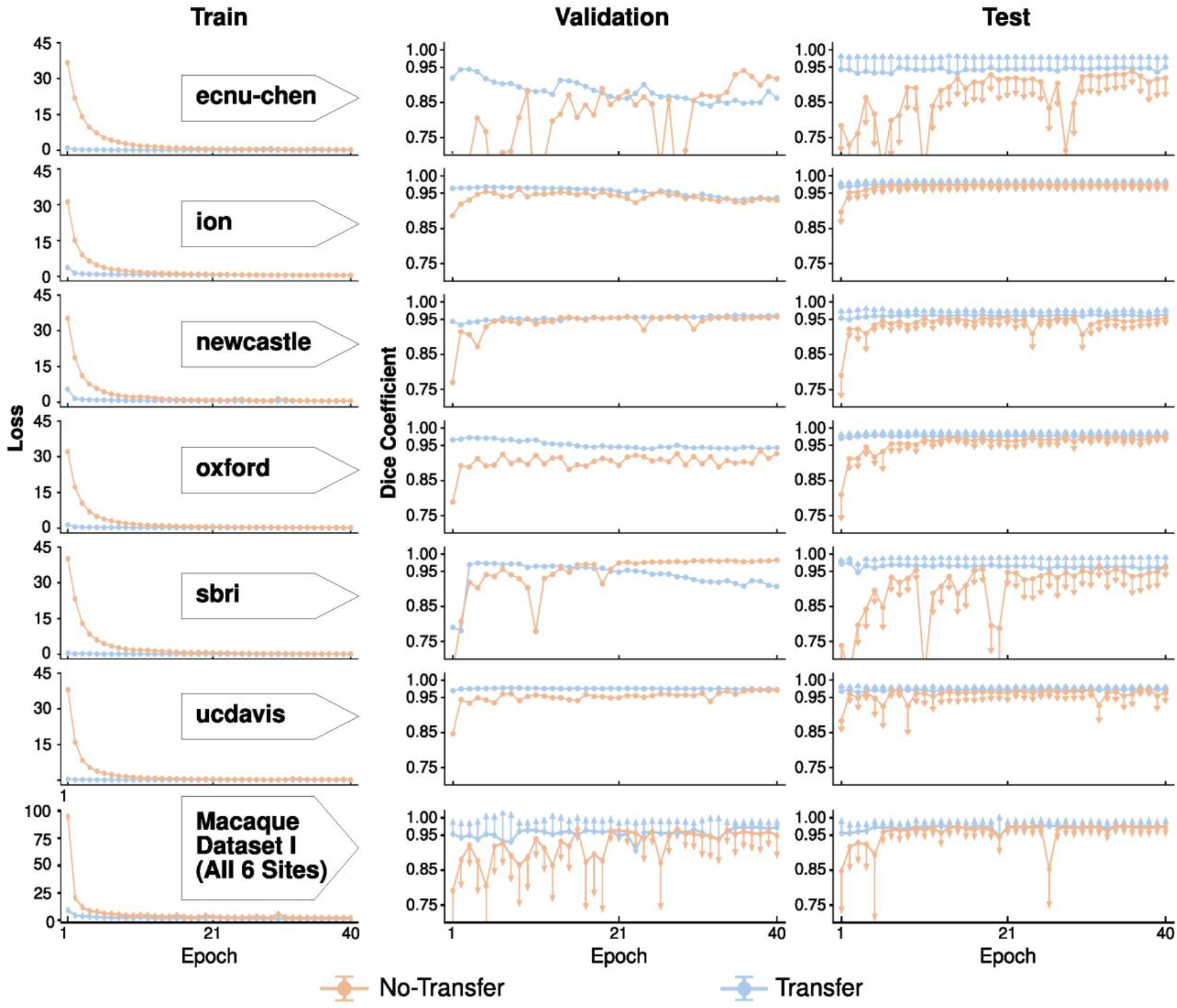
Comparison of skull-stripping performance of the U-Net models with and without transfer-learning on Macaque Dataset I. The models with transfer-learning (blue curve) outperform the one without (orange curve) across each of six sites (row 1-6), as well as the merged sample (the last row). The loss (i.e. the sum of cross-entropy, first column) between the predicted mask and ground truth mask on the train set converges faster for the model with transfer-learning. Similarly, models with transfer-learning show higher Dice coefficients with lower variation in the validation and testing sets than models without transfer-learning. Of note, one-side error bars (standard deviation) were used to avoid dense overlaps between the two models.

We also computed and evaluated the model with and without transfer-learning based on the merged training samples across six sites (N=12). Similarly, the transfer-learning model (referred to as U-Net T12 model) showed lower loss and higher Dice coefficients across 40 epochs than the model without transfer-learning (Fig. 3, last row); its best performance epoch (i.e. the 37th epoch) was selected as our macaque transfer-learning model (i.e. U-Net T12).

### 3.3. Comparison Between the U-Net Model and Traditional Approaches

Recognizing that the model with transfer-learning is superior to the one without, we then updated the transfer-learning model (i.e. U-Net T12) using Macaque Dataset I (N=12) & II (N=7) to generate our final generalized model (U-Net 12+7 model). Here, we evaluated the performance of the U-Net T12 model and the U-Net 12+7 model in comparison to other traditional brain extraction approaches. Brain masks from the two U-Net models showed significantly higher Dice coefficients than those from the traditional pipelines (Fig. 4, *F*=30.164, *p*<10^−29^ repeated ANOVA, all post-hoc p<0.05). Skull-stripping using the U-Net models was successful (Dice>0.95) for all of the testing macaques (N=30) across six sites. Notably, the U-Net 12+7 model showed relatively higher Dice coefficients than the U-Net T12 model (Fig. 4, *paired-t*=3.62, *p*=0.001), though the additional training samples used in U-Net 12+7 model were not included in the sites where the testing samples were selected from. This indicated the generalizability of model-upgrading across sites. At the voxel level, both the U-Net T12 and ‘12+7’ models exhibited fewer false negatives and false positives than traditional pipelines (Fig. 5). The U-Net T12 model showed slightly more false positives than false negatives, which indicated that the model tends to include a few non-brain voxels on the edge of the brain mask rather than miss the brain tissue. Overall, these results demonstrated the feasibility of transfer-learning and model-upgrading using small training samples.

**Figure 4.**
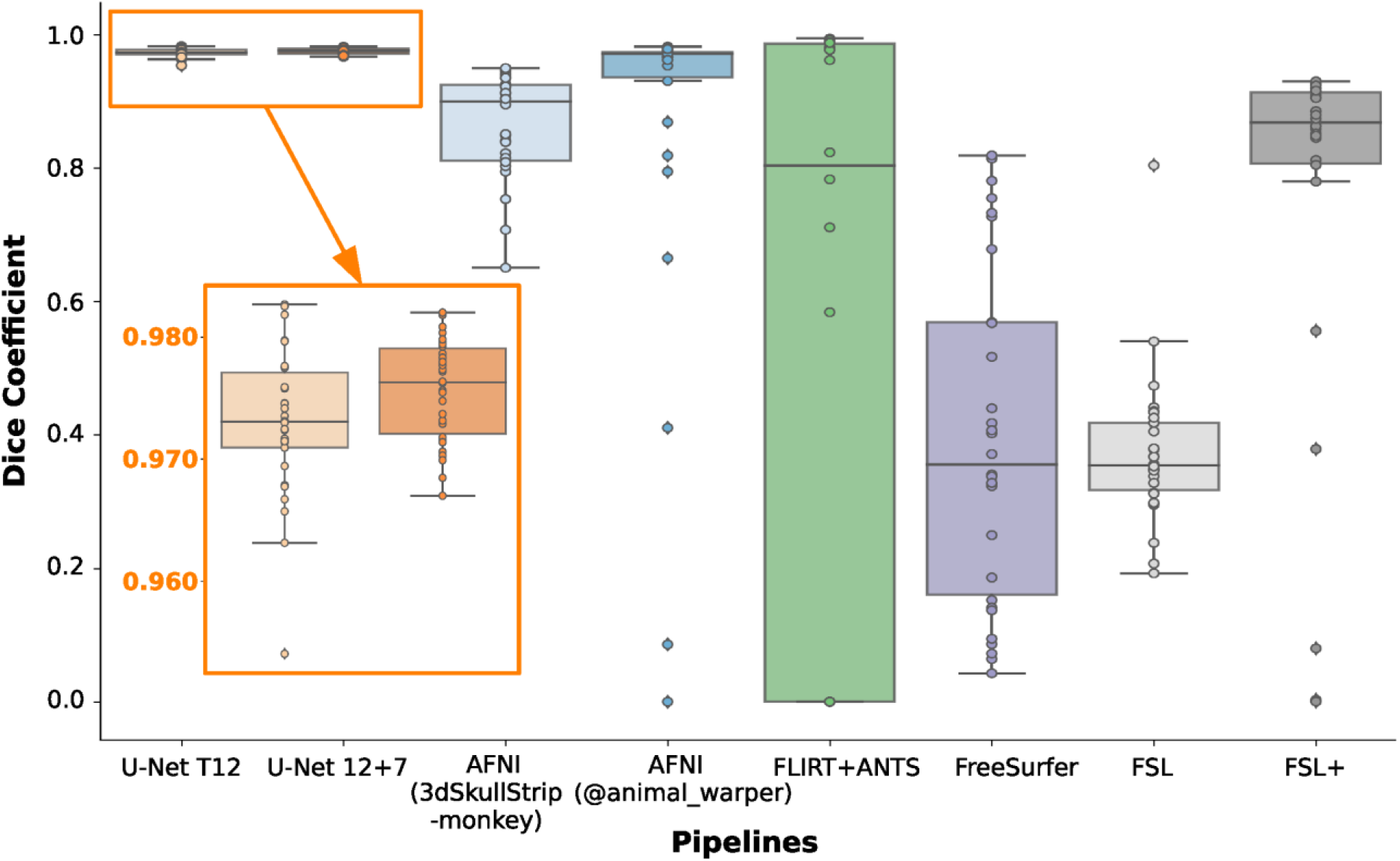
Performance of the U-Net models and traditional approaches. The boxplot shows the Dice coefficients in the testing datasets (N=30) of Macaque Dataset I across brain extraction approaches including 1) the transfer-learning model (i.e. U-Net T12), 2) generalized model (i.e. U-Net 12+7), 3) AFNI 3dSkullStrip command with ‘-monkey’ option, 4) AFNI @animal_warper pipeline, 5) template-driven FLIRT+ANTS pipeline, 6) FreeSurfer HWA approach, 7) FSL BET approach, and 8) FSL BET approach with customized options (FSL+).

**Figure 5.**
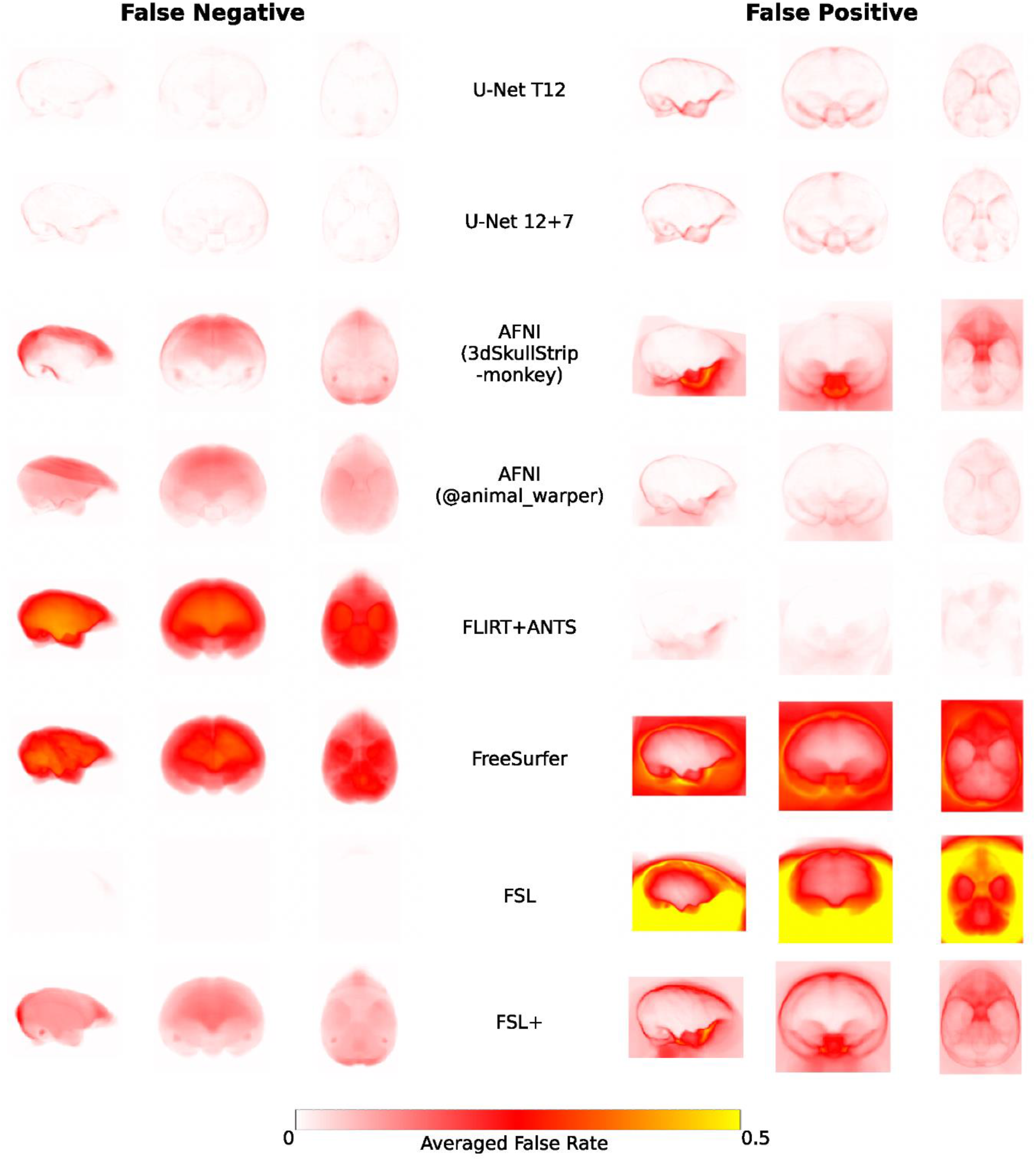
The averaged voxel-wise false-negative and false-positive rates of the U-Net models and traditional approaches in the testing datasets of Macaque Dataset I. The false-negative rate examines where the predicted mask falsely misses the brain tissue in the brain (left column) while the false-positive rate examines where the predicted mask falsely includes the non-brain tissue. Of note, FSL BET tends to falsely include large amounts of non-brain tissue in the brain mask, thus false-negative rates are close to zeros.

Comparing amongst the traditional approaches, the template-driven approaches (i.e. AFNI @animal_warper and Flirt+ANTS), and AFNI 3dSkullStrip with parameters customized for NHP data (3dSkullStrip -monkey) showed better performance than FSL and FreeSurfer (Fig. 4–5). FSL performed better with the radius setting (Fig 4–5, FSL vs FSL+), though still not particularly accurately (Dice<0.928). Both template-driven approaches appear to be more conservative and have missed the brain tissue (low false positives and high false negatives). AFNI 3dSkullStrip missed identifying the brain in the superior regions (higher false negatives on the top) and falsely included non-brain voxels in the bottom (Fig. 5).

### 3.4. Generalizability of U-Net model and Skull-Stripping PRIME-DE Samples

To evaluate the generalizability of the U-Net model, we applied the U-Net T12 and 12+7 models to all the other macaque samples (i.e. T1w images) contributed to PRIME-DE from differing research sites. The results of brain masks were visually reviewed by three experts and rated into four grades (good, fair, poor, and bad). Good and fair ratings were considered to be successes, while poor and bad ratings were failures. We also used the traditional pipelines and similarly rated their brain mask results. The performance of different pipelines for each macaque is shown on github repository: https://github.com/HumanBrainED/NHP-BrainExtraction. Figure 6 shows the proportion of macaques with good, fair, poor, and bad skull-stripping masks for each of the 20 sites, for each of the five approaches. Again, the U-Net models outperformed the traditional approaches for most sites. In particular, the final U-Net 12+7 model showed the best performance across macaque samples (success rate=90.4%). All macaque samples (N=123) were successfully skull-stripped across the twelve sites with thirteen exceptions. This result demonstrated the generalizability of the U-Net 12+7 model across sites. Of note, all crab-eating macaques *(M. fascicularis*, N=12) were successfully skull-stripped using the U-Net 12+7 model, though the training samples only contained the rhesus macaques (*M. mulatta*). No significant difference was observed in age and sex between the successes sand failures. In addition, four out of five sites with anisotropic data also succeeded (sbri: 0.6×1.2×0.6 mm, nki: 0.50×0.55×0.55 mm, ecnu and ecnu-chen: 0.75×0.75×0.8 mm75×0.75×0.8mm, NIMH: 1.5×0.5×0.5mm). These findings suggested the generalizability and relative robustness of the U-Net model across macaque species, age, sex and whether the voxels were anisotropic or isotropic.

**Figure 6.**
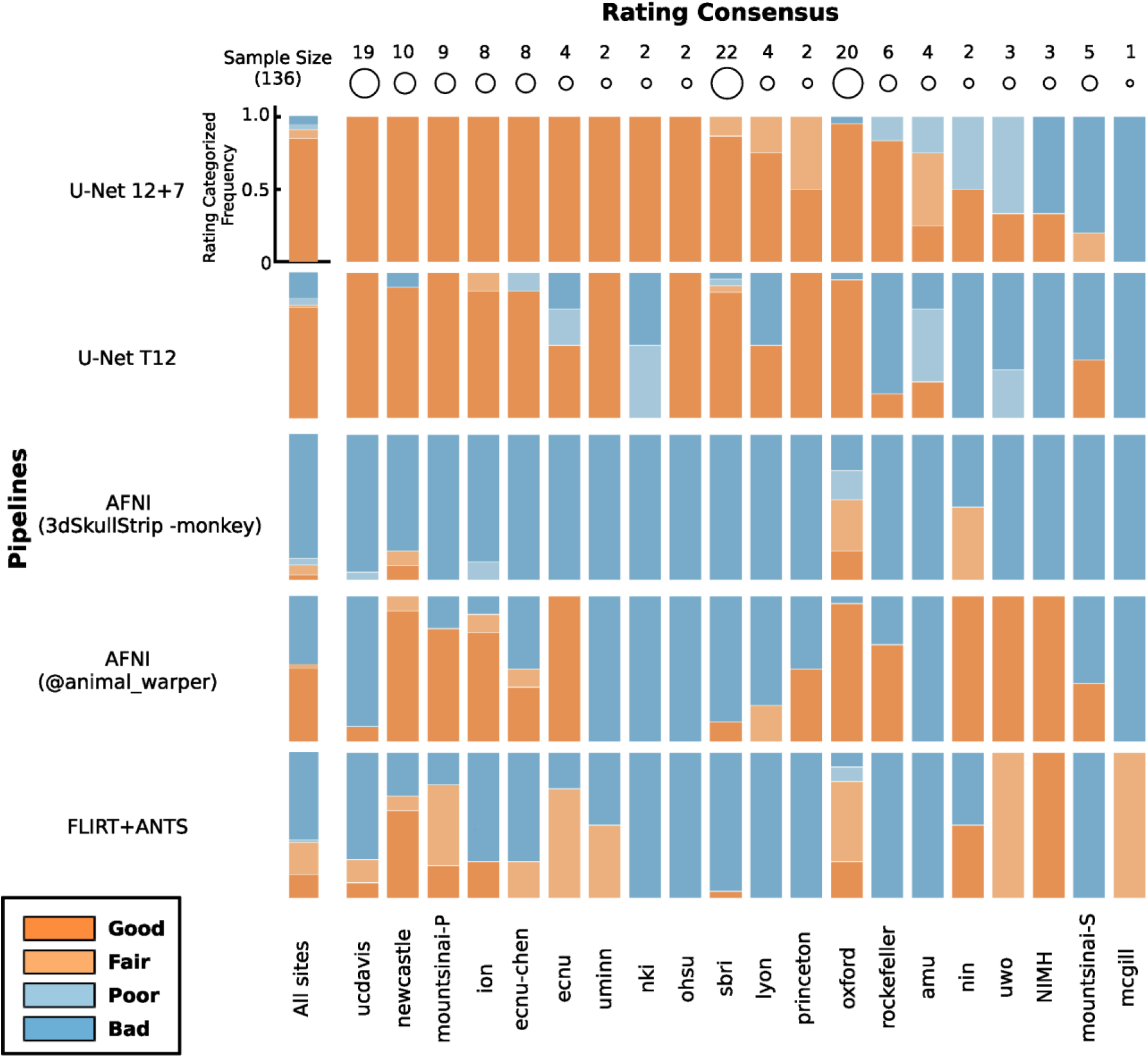
Expert rating consensus of brain extraction performance across the U-Net models and traditional approaches for the PRIME-DE datasets. The stacked bar plots show the proportion of macaques with good, fair, poor, and bad skull-stripping masks on T1w images for each of 20 sites in PRIME-DE. Of note, the FSL BET customized pipeline only succeeded for 5 macaques. FreeSurfer HWA and FSL BET default pipelines failed to obtain fair/good brain masks, and thus their ratings were not displayed. The U-Net 12+7 model (the first row) shows the highest rating across pipelines.

Among intensity-based approaches, AFNI 3dSkullstrip showed better performance (successful rate=10.3%) than FSL and FreeSurfer. FSL and FreeSurfer default pipelines failed in almost all the cases except FSL with customized options (successful rate=3.7%). In particular, massively incorrect results were observed (see visual inspection figures: https://github.com/HumanBrainED/NHP-BrainExtraction). This is because the intensity-based approaches heavily depend on the data quality. Specifically, these approaches perform brain extraction by identifying the center of the brain and expand the brain outline outward till the intensity drops (at the cerebrospinal fluid [CSF] or skull). As such, it is crucial to set the brain/head radius as the NHP has a smaller brain size than the human, recognize the center of the brain for the input image, and have a sufficient intensity contrast between brain and CSF/skull to identify the brain boundary. However, as shown in Fig 1, those conditions in NHP data are usually not satisfied. Thus, most of the samples failed with FSL, FreeSurfer, or even AFNI 3dSkullstrip with ‘-monkey’ option.

We also noticed that template-driven approaches performed well in some cases but failed in others (success rate: AFNI @animal_warper=52.9%, Flirt+ANTS=38.2%). Visual inspection of the intermediate outputs from AFNI @animal_warper and Flirt+ANTS pipelines showed that all successful cases had the first linear alignment of the individual’s head to the template head somewhat close. The failures mostly occurred in the first linear registration step; 79.4% of 63 skull-stripping failures using @animal_warper and 89.3% of the 84 failures using the Flirt+ANTS pipeline had failed in the first linear registration step. We also noticed that, for some cases, the brain mask generated by AFNI @animal_warper had a straight cut-off at the top of the brain (Fig.(Fig S5A). This might be caused by the displaced center of the brain within these T1w image. As a first step, @animal_warper aligns the centers of the input and template volumes. If the brain is far from the center of the image, this first pass at alignment with the template may be too far off for subsequent alignment steps to succeed. We found that re-centering the head to the center of the image could fix the straight cut-offs, though did not necessarily produce a good brain mask (Fig. S5B). We further cropped the neck regions by zeroing out the bottom slices using the U-Net output as a prior. Specifically, we zeroed the bottom k slices along the z-axis. Here *k* was determined by *N-m-p*, where N = the total number of slices of the image along the z-axis, m = the total number of non-zeros slices of the U-Net mask, p = the number of slides from the top slice of the image to the first non-zero slice of U-Net mask along the z-axis. After the recentering and cropping, we found that @animal_warper turned most of the failed cases (66.7% of 63 failures) into successes (Fig S5C). For some cases, however, inaccuracies remained (Fig. S5D).

U-Net successfully skull-stripped 123 out of 136 macaques. Among the 13 failures in the U-Net 12+7 model, 8 macaques failed because their T1w images were substantially different from the training samples (e.g., highly inhomogeneous intensity, very different field of view, Fig. S6A). For the other 5 failed cases, the U-Net 12+7 model still enabled identifying most of the brain with only minor inaccuracies at the edge of the brain (Fig. S6A). When we used the U-Net output mask to perform an initial brain extraction and estimated the affine transformation from the initial skull-stripped brain to the template brain, the template-driven approaches (AFNI @animal_warper and ANTS) turned seven failed cases into successes (Fig S6C). As the performance of the deep-learning approach heavily depends on the training set, we also updated the U-Net 12+7 model using site-specific training data and showed the improvement across macaques (Fig S6B). We have selected the best of the successful brain masks (Fig S6D) for each macaque and shared them on PRIME-RE (Messinger et al., 2021).

### 3.5. Application of U-Net Model-Upgrading for an External Dataset and MP2RAGE Images

Here we used a large external dataset (N=454, UW-Madison dataset with manually drawn whole-brain masks, details described in previous studies) to evaluate the utility of our U-Net -based approach (Fox et al., 2018, 2015; Oler et al., 2010). First, we directly applied the model-prediction module using our U-Net 12+7 model to extract the brain masks for 40 macaques and found relatively good results (Dice=0.923+/−0.025). We further used the U-Net 12+7 model as the pre-trained model and upgraded the U-Net model on these 40 macaques for 10 epochs. This procedure took about 90 min. The upgraded model showed substantial improvement and successfully skull-stripped all remaining macaques (N=414, All Dice>0.95, Mean=0.977±0.005). We further challenged our U-Net model-upgrading module with MP2RAGE images collected from the UWO site (N=8) in PRIME-DE, which have different intensity profiles with opposite contrast in gray matter and white matter (Fig. S7). By upgrading the pre-trained U-Net 12+7 model with three MP2RAGE brain masks, the U-Net model enabled skull-stripping on the rest of the hold out MP2RAGE images (N=5).

### 3.6 Applications of U-Net Transfer-learning Model to Chimpanzee, Marmoset and Pig Samples

To demonstrate the feasibility of applying our U-Net tool in other primates, and other mammals, we included chimpanzee data (N=1) from a repository of previously collected scans (the National Chimpanzee Brain Resource; https://www.chimpanzeebrain.org), the common marmoset template data (N=1) from the Riken marmoset atlas (https://brainatlas.brain.riken.jp/marmoset_html), a common marmoset dataset (N=5) from the coauthor SS, and a pig dataset (N=5) (Benn et al., 2020). We directly applied the U-Net 12+7 model to the chimpanzee data (Figure 7A). Interestingly, the U-Net 12+7 macaque model performed relatively well in chimpanzees. We also applied the U-Net 12+7 macaque model directly to the marmoset data. The result was fairly good for the template data (Fig 7B) but failed in the individual data with highly site-specific inhomogeneous noise (Fig 7C). We further updated our 12+7 macaque model using one manually edited marmoset. The updated model succeeded in the rest of the 4 marmosets from the same site (Fig. 7D). We also tested whether our U-Net tool is capable of transferring the macaque model to other mammals. With the addition of training data from 3 pigs, the transferred model achieved a Dice coefficient ≥ 0.93 in the remaining 2 pigs (Fig. 7E).

**Figure 7.**
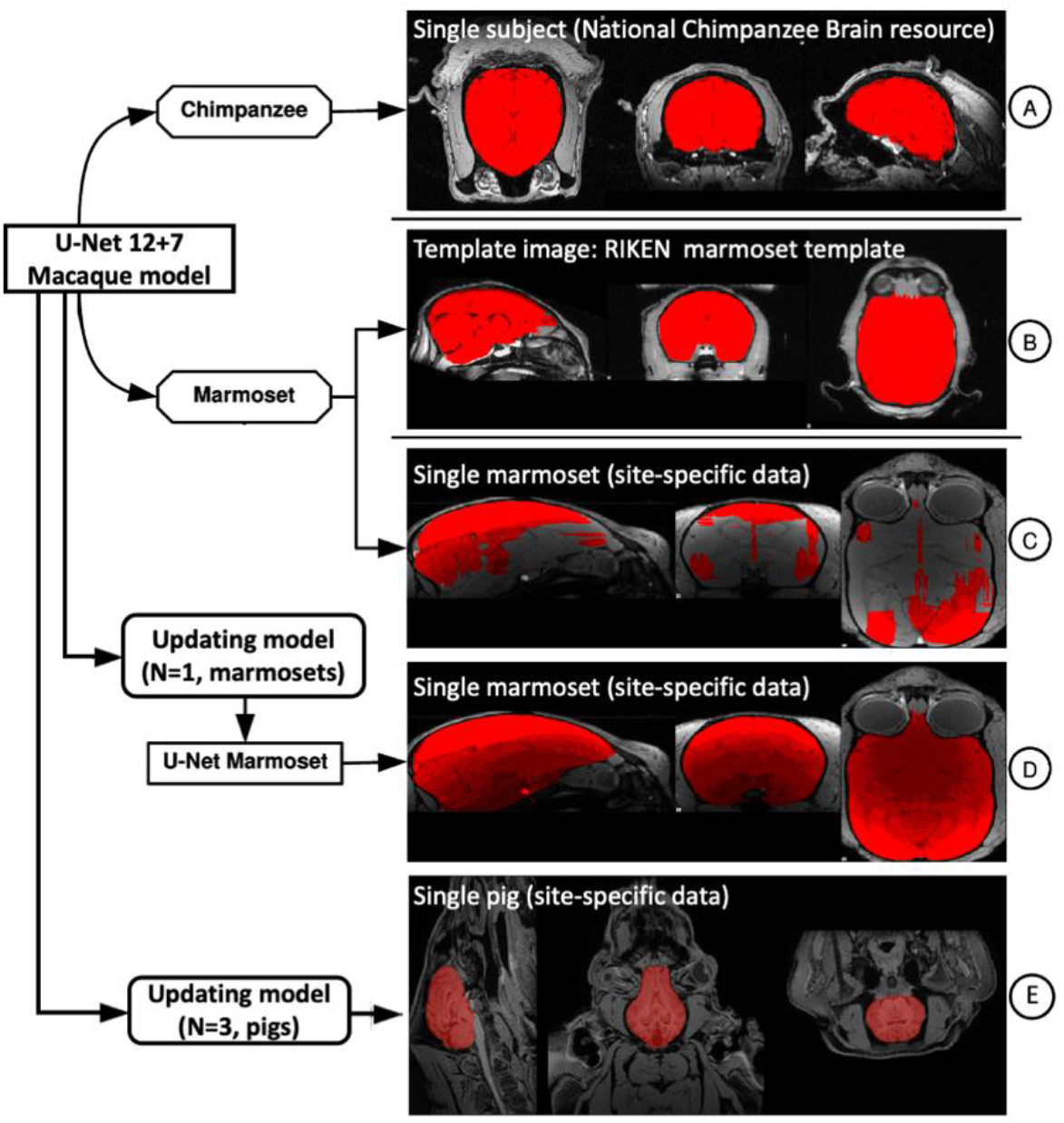
The applications of the U-Net tool in the chimpanzee, marmoset, and pig datasets.

## 4. Discussion

The present work demonstrated the feasibility of developing a generalizable brain extraction model for NHP imaging through transfer learning. Central to the success of our effort was our leveraging of human data as a base training set, upon which additional learning for NHP and site-related variations could be readily achieved. We employed the heterogenous, multi-site PRIME-DE resource to evaluate the effectiveness of our framework, finding that the transfer-learning U-Net model identified macaque brain tissue with higher accuracy than traditional methods and also proved more generalizable across data collections. Notably, the transfer-learning U-Net model provides a notably faster solution (approximately 1-10 min) than the next best performing algorithms, which tend to rely on template-based strategies and require over an hour. We have released our model and code, including the utilities for skull-stripping, model-training, transfer-learning, as well as model-updating modules (https://github.com/HumanBrainED/NHP-BrainExtraction). Additionally, we created and shared the skull-stripped repository of 136 macaque monkeys to facilitate the large-scale macaque MRI imaging for the PRIME-DE and NHP community (Messinger et al., 2021; Milham et al., 2018).

*A priori*, the major roadblocks that one would anticipate for the application of deep learning in NHP imaging are the small sample sizes and variations in imaging protocols (Autio et al., 2020b; Milham et al., 2018). In part, this is a reflection of the experiences of the human imaging community, where large sample sizes have been required to accurately segment the brain using deep learning (Henschel et al., 2020; Kleesiek et al., 2016). The superiority of the transfer learning models in the present work emphasizes the unique advantages of transfer-learning in overcoming such challenges for nonhuman primate imaging and offers a model that may be considered in future efforts to overcome similar obstacles in other imaging populations (e.g., macaques with surgical implants and/or in-dwelling electrodes, brain extraction in other species, pathologic models, early development, aging) (Pontes-Filho et al., 2019; S. Roy et al., 2018; Salehi et al., 2018). The success of the transfer-learning model emphasizes the similarity of the general structure of brain tissue in both species (e.g. gray matter, white matter) - despite the anatomical differences between human and macaque heads (e.g. head size, muscular tissue surrounding the skull, skull thickness, etc.) (Yosinski et al., 2014). Aside from accuracy, the transfer-learning model also converged faster (leveling off after 2 epochs) and yielded more stable results across epochs, even though the pre-trained model was established using a different species. Notably, our human-to-macaque model also enables skull-stripping the chimpanzee data though no chimpanzee training samples were used. Moreover, the smaller NHP (e.g. marmoset) and other mammals (e.g. pig) could also benefit from the transfer-learning model from the human-to-macaque and only required adding a small training sample to update the model. These findings suggest the utility of transfer-learning in other animal studies and further brain tissue segmentation.

The success of transfer learning in the present work may signal the ability to use smaller samples than previously employed for human imaging studies (Ghafoorian et al., 2017). However, this is not necessarily the case, as this may instead reflect that the folded surface of the macaque is much less complex than that of humans. The macaque central sulcus is less meandering on the lateral parietal lobe (Hopkins et al., 2014). There is only one superior temporal sulcus in the temporal lobe and two less curved sulci (i.e. rectus, arcuate) in the frontal lobe, such that there is a relatively smooth surface edges for brain extraction (Bogart et al., 2012; Hopkins, 2018). In addition, such folded and meandering brain morphology is substantially more similar across individual macaques than humans (Croxson et al., 2018). As such, small training samples suffice for the macaque model compared to the human. Future work in the human imaging community would benefit from a systematic examination of minimal sample sizes needed for successful training and generalization.

Beyond transfer across species, a key finding of the present work is the ability to improve model generalizability across independent imaging sites relative to traditional methods. By further upgrading the pre-trained transfer-learning model based on the secondary training samples across multiple datasets, the upgraded U-Net model has improved the brain extraction performance and showed a higher success rate than the traditional methods across multiple sites. Of note, the upgraded model enabled successful skull-stripping of datasets acquired from three additional sites that were not included in any training sites. This demonstrates the out-of-site generalizability of upgrading the pre-trained U-Net model across sites. More impressively, we found that the model-upgrading module offers a solution of generalizing the pre-trained model to other modalities (i.e. MP2RAGE), which is usually difficult to achieve with traditional methods. These findings highlight the important role of pre-trained models in brain extraction for small samples, regardless of site (across or within sites), modality (MPRAGE or MP2RAGE), and species (across or within species). Further improvement for the user-specific dataset can be achieved by adding small samples (N≥2) using our U-Net result as the pre-trained model.

It is worth noting that the U-Net model we used in the current study is a 2D convolutional neural network. Although the 3D model usually tends to have higher accuracy (Hwang et al., 2019), the 2D model has a smaller network size, much less memory cost, and requires less computational time. More importantly, the 2D model is more accessible in general computational platforms for users without large amounts of video memory, which can be costly. In addition, unlike the single slice 2D model, our 2D model leveraged the local interslice features by using each slice and its neighboring slices (i.e. 3 slices in total as opposed to just one) as the input image in the first layer. We also resampled the slices in axial, sagittal, and coronal planes and combined the predictions from all three planes into a final probability map (Lyksborg et al., 2015). By doing so, we effectively tripled the number of training slices, which is especially useful for optimizing prediction when training sample sizes are limited (i.e. data augmentation technique). Of note, we opted for the U-Net model as the U-Net-like methods appear to be preferred for a broad biomedical image application across a variety of segmentation designs (e.g. identifying heart, cell, tumor, vessel etc.) (Isensee et al., 2021). For further improvement in a specific application, besides optimizing the specific architecture of neural networks, adding training data and optimizing the preprocessing step (e.g. denoising, bias correction, data harmonization etc.) are still suggested. For example, a conditional random field can be considered in the prediction layer for further refining the weights in the tissue prediction (Chen et al., 2019; Zhao et al., 2018). Beyond improving brain extraction, future efforts may place a greater focus on tissue classification (e.g. GM, WM, subcortical structures). Central to such efforts will be the sharing and amassing of manually segmented brain images, to which 3D CNNs can be applied.

To promote pipeline development in the NHP field, we have released the skull-stripped brain masks, our generalized model, and code via the PRIMate Resource Exchange platform (PRIME-RE: https://prime-re.github.io) (Messinger et al., 2021). Researchers can access the code and perform the brain extraction on their own macaque datasets. The model-prediction for a dataset takes about 20 seconds on a GTX1070 GPU with 700MB GPU memory or 2-10 min on a single CPU core. We also included the model-building and model-upgrading modules in the code, which has been implemented in a recent version of a Configurable Pipeline for the Analysis of Connectomes (C-PAC v1.6: https://github.com/FCP-INDI/C-PAC/releases/tag/v1.6.0). When the results from the current model need improvement, users can upgrade the U-Net using our current generalized model as a pre-trained model and expect a stabilized solution after several training epochs (N<10 epochs) without validation datasets. The model-upgrading module takes about 1-5 hours on a single CPU, or 15-20 min on a GPU. In addition, we released our manually skull-stripped masks (40 macaques across 7 sites) which can be used as ‘gold standards’ for other deep-learning algorithms. Researchers are encouraged to manually refine brain masks to further improve the training datasets (e.g. NMT v2.0) for deep-learning approaches (Jung et al., 2020; Seidlitz et al., 2018). Additionally, we included the successful brain mask outputs to facilitate the further preprocessing analysis of PRIME-DE data. For a new macaque dataset, we recommend one to:

1. Use the U-Net 12+7 model first.
2. If the U-Net 12+7 model fails with minor inaccuracies, consider using the U-Net output to perform an initial linear alignment and apply AFNI @animal_warper or ANTs.
3. If the U-Net 12+7 model fails with high inaccuracies, use AFNI @animal_warper as the alternative. Of note, we recommend to recenter and crop the image before applying @animal_warper.
4. If both the U-Net 12+7 modal and template-based approaches fail, manually edit a few (N>=1) datasets and update the U-Net model using the 12+7 model as a prior.

There are some limitations in our studies. First, although the final U-Net model showed better generalizability across research centers than traditional approaches, it is unable to accurately skull strip all macaque datasets (success rate: 90.4%). Of note, in failed cases, it is possible to use the output of the U-Net model as the initial brain mask to recenter and crop the image, and then create a prior linear transformation for traditional template-driven approaches - a process that can turn most of the failed cases into successes. Second, the U-Net model requires denoising and bias correction using traditional approaches prior to the model prediction. Future work may consider leaving this image noise and the bias field information in the image during training of the network to simplify the processing steps and possibly improve performance.

## 5. Conclusion

In the present work, we proposed and evaluated a fast and stable U-Net based pipeline for brain extraction that exhibited performance superior to traditional approaches in a heterogenous, multisite NHP dataset. We have released the code for brain mask prediction, model-building, and model-updating, as well as macaque brain masks of PRIME-DE data. We hope this open repository of code and brain masks can promote pipeline development in the NHP imaging field and accelerate the pace of investigations.

## Acknowledgement

This work was supported by gifts from Joseph P. Healey, Phyllis Green, and Randolph Cowen to the Child Mind Institute, fundings from National Institutes of Health (NIH BRAIN Initiative Grant R01-MH111439 to CES and MPM, P50-MH109429 to CES, R24MH114806 to RCC and MPM, and R01MH120482 to TX). This research was supported (in part) by the Intramural Research Program of the NIMH and includes the relevant Annual Report number in the following format (ZIAMH002918 to AM). This research was also partially supported by funding from NIMH R01MH121735 to ASF, the California National Primate Research Center (P51OD011107 to ASF), NIH R01MH101555 to RCC, R01MH081884 to NHK, R01MH046729 to NHK, P50MH084051 to NHK, National Nature Science Foundation of China (81571300, 81527901, 31771174 to ZW), NIH BRAIN Initiative Grant RF1MH117040 to BER, Wellcome Trust Investigator Award (108089/Z/15/Z) and Medical Research Council Programmer Grant MR/ M023990/1. We also thank the PRIMate-Data/Resource Exchange consortium (PRIME-DRE).

## Supplementary Table

**Table S1.** The Pipelines, command options, approximate processing time, and successful rate estimated using 136 macaque samples from PRIME-DE.

| Pipeline | Options | Processing times | Dice coef (N=30, testing set) | Success rate (N=136) |
| --- | --- | --- | --- | --- |
| AFNI<br>3dSkullstrip | 3dSkullstrip -push_to_edge -monkey -shrink_fac 0.5 | ~1-2 min | 0.841±0.134 | 10.3% |
| FSL | bet -f 0.3 | ~30 sec<br>- 1 min | 0.374±0.110 | 0% |
| FSL+ | bet -f 0.3 -g -0.5 -r 35 | ~30 sec<br>- 1 min | 0.739±0.308 | 3.7% |
| FreeSurfer | mri_watershed default | ~30 sec<br>- 1 min | 0.396±0.250 | 0% |
| FreeSurfer+ | mri_watershed -T1 -atlas -r 30/voxel_size | ~30 sec<br>- 1 min | 0.231±0.210 | 0% |
| Flirt+ANTS | flirt -dof 6<br>ANTS -t SyN[0.25] -r Gauss[3,0] -i 60x50x20 --use-Histogram-Matching --number-of-affine-iterations 10000x10000x10000x10000x10000 -MI-option 32x16000 | ~30 min –<br>1 hour | 0.556±0.471 | 38.2% |
| AFNI<br>@animal_warper | @animal_warper<br>Template: NMT 0.5mm | ~2 hours | 0.865±0.254 | 52.9%<br>(84.6% *) |
| U-Net prediction | muSkullStrip.py | ~2 min | 0.98±0.004 | 90.4% |
| U-Net training | trainSs_UNet.py -init -epoch 10 (default=40) Training set (N=2), Validation set (N=0) | ~10-15 min (with GPU)<br>~1-2 hours | n.a | n.a |

\* The success rate was improved after recentering and cropping the input image using the U-Net output as an initial brain mask for @animal\_warper.

## Supplementary Figures

**Figure S1.**
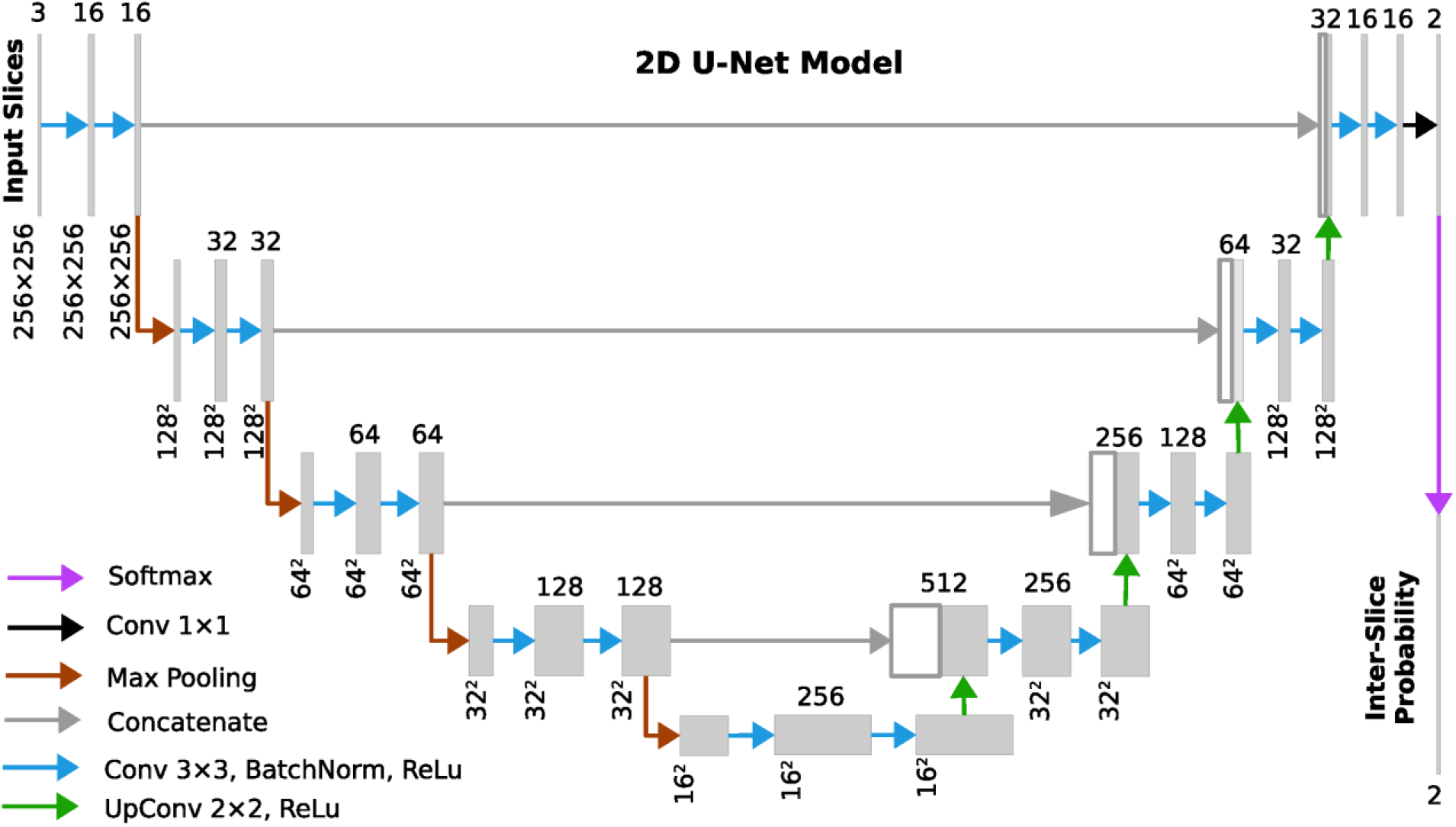
Outline of the architecture of 2D U-Net.

**Figure S2.**
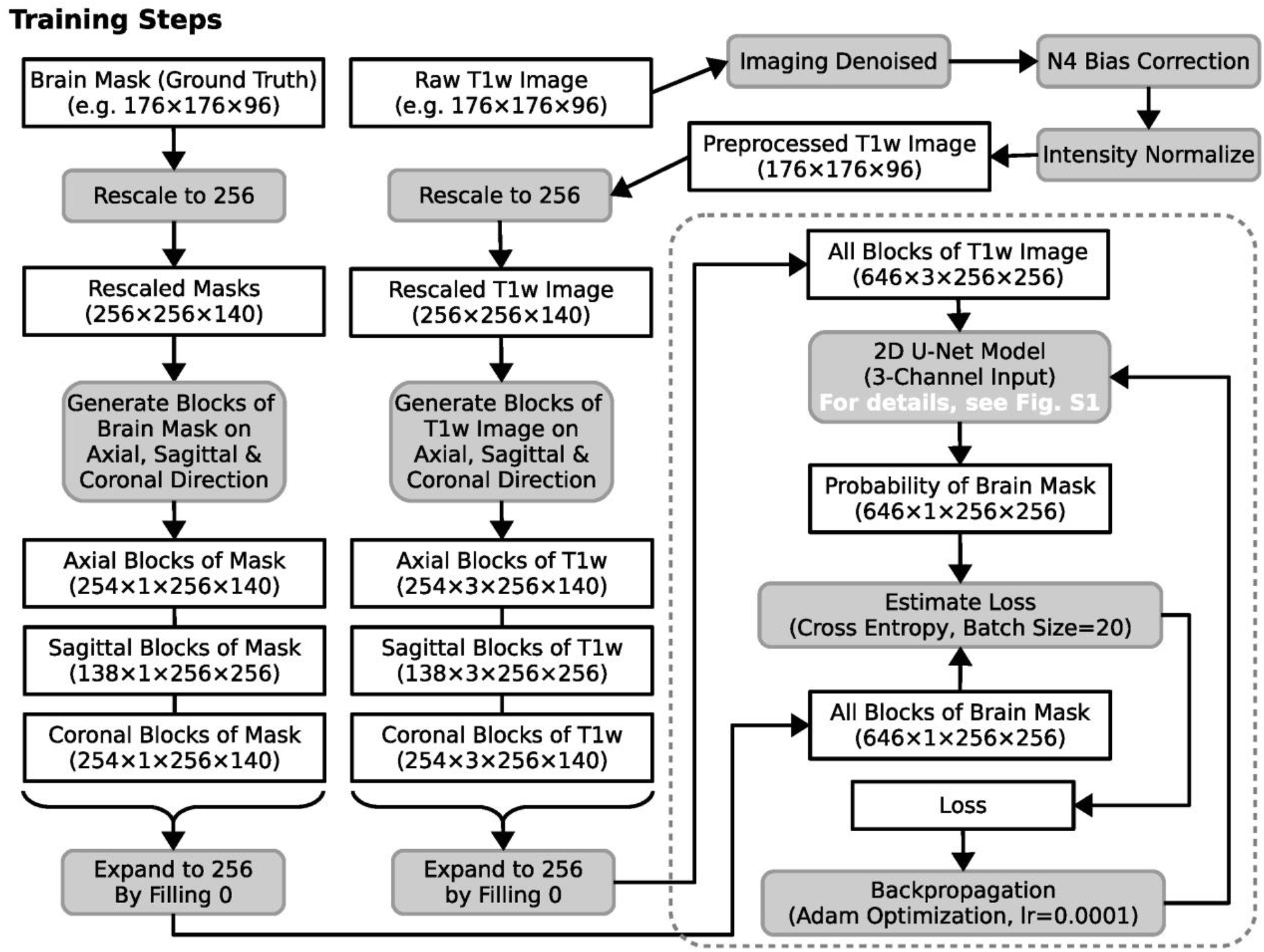
U-Net training procedure for each 3D image.

**Figure S3.**
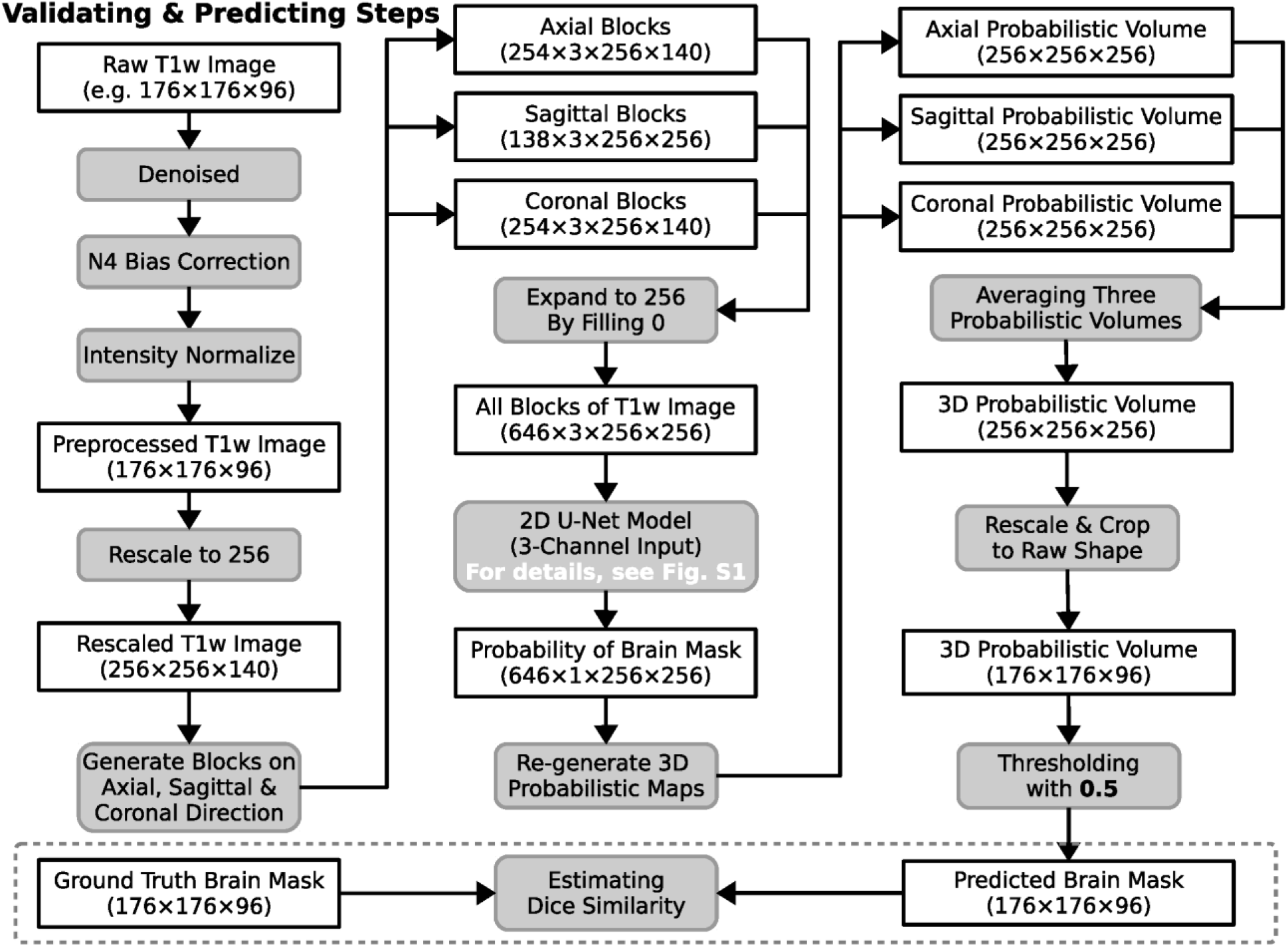
U-Net validating and prediction procedure for each 3D image.

**Figure S4.**
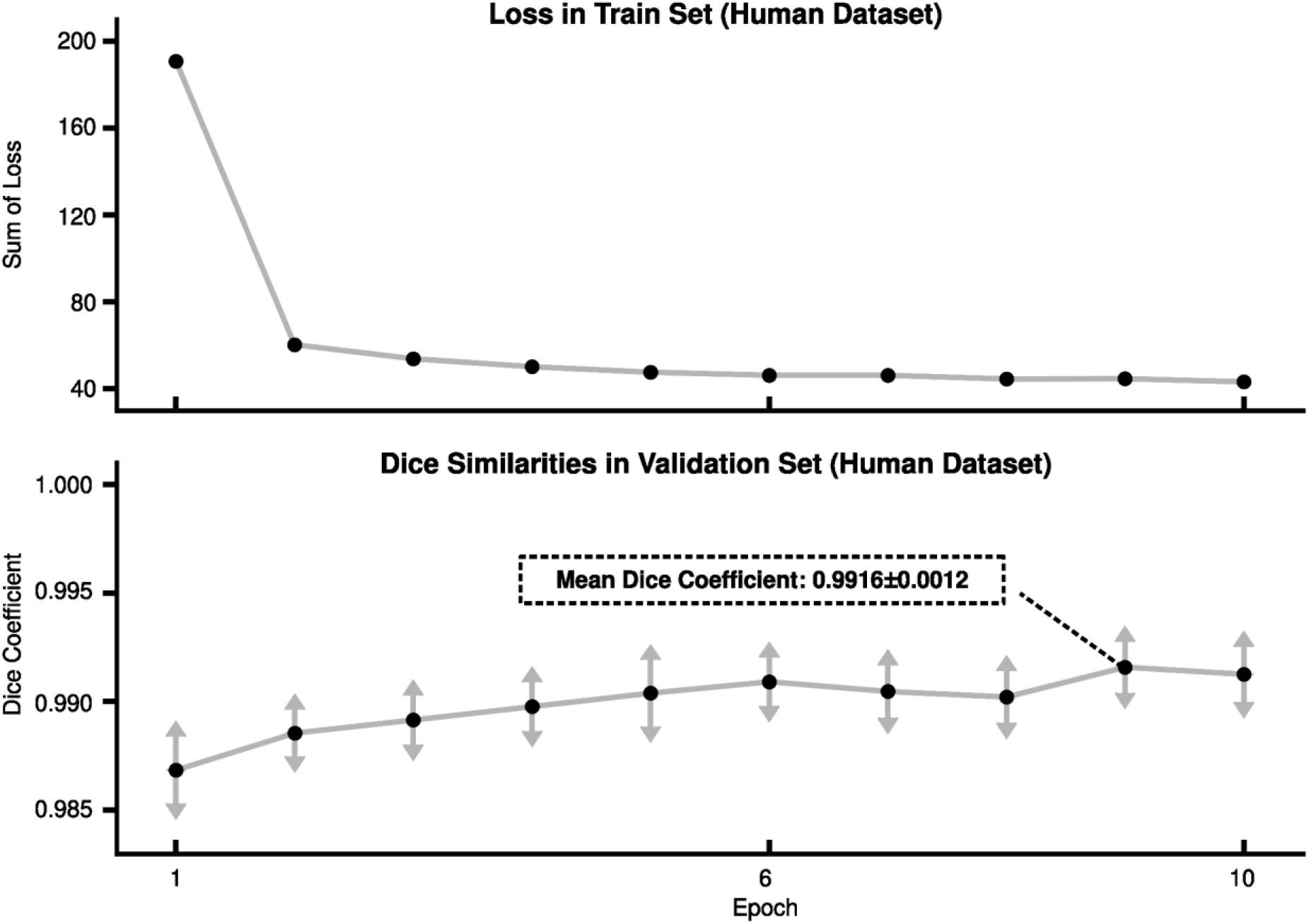
The loss and Dice coefficient of the U-Net model for each training epoch on the human dataset. The mean and standard deviation (arrows) across validation samples was calculated for each epoch.

**Figure S5.**
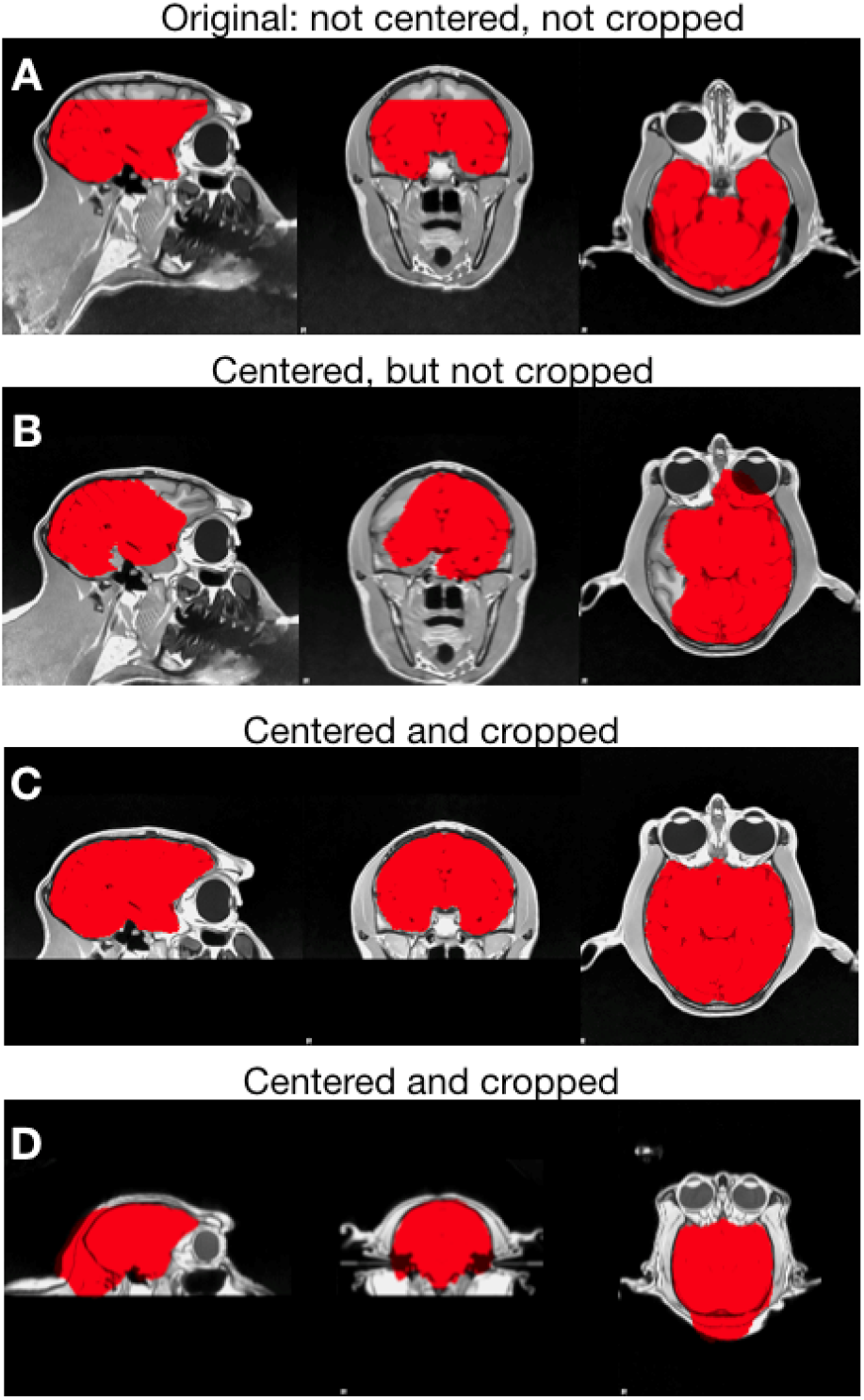
Examples of AFNI @animal_warper results for different inputs. Brain masks for (A) the original T1 image (not centered or cropped), (B) the images centered (the brain is placed in the middle) but not cropped (the neck is included), (C) The input image is centered and cropped, resulting in an @animal_warpper success. (D) The input image is centered and cropped, yet @animal_warpper still failed.

**Figure S6.**
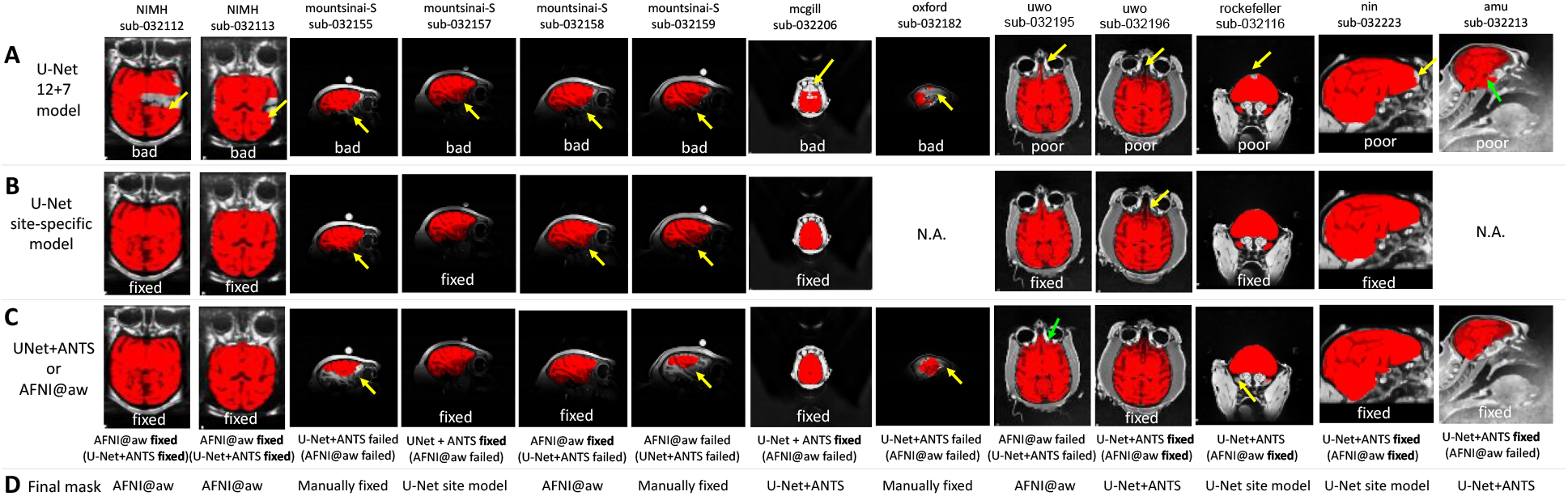
Visual inspections of all failed cases in the U-Net 12+7 model. **A)** brain masks and rating using the U-Net 12+7 model. **B)** brain masks and rating using updated the U-Net site-specific model. **C)** brain masks obtained using either the AFNI @animal_warper or U-Net+ANTs template-driven approach. The better of the resultant masks is shown. The tool used and the outcome obtained are indicated below the image. The alternative method and outcome are reported in parentheses. **D)** the method producing the best mask selected for each macaque.

**Figure S7.**
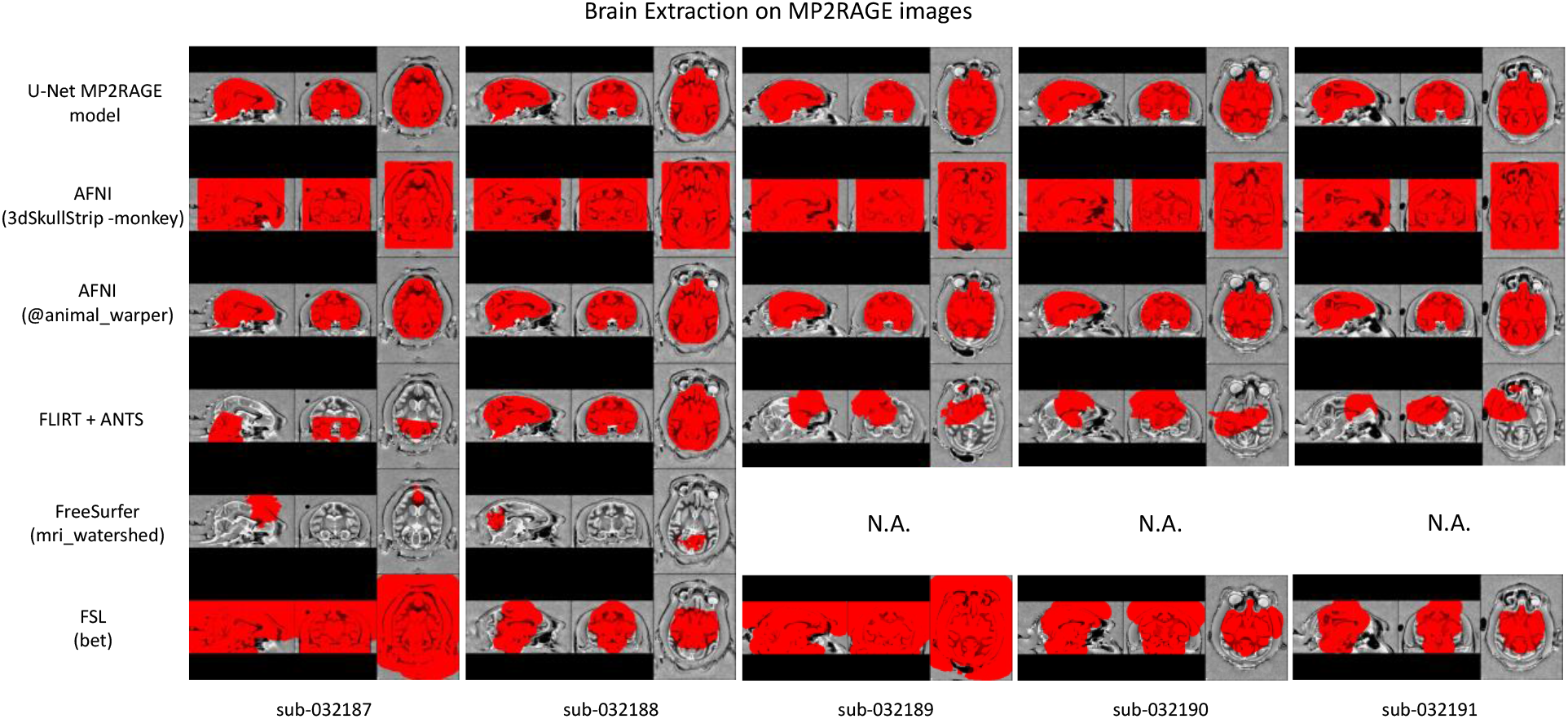
Skullstripping results of U-Net MP2RAGE model and traditional pipelines for MP2RAGE dataset from PRIME-DE (site-uwo).

## Reference

Acosta-Cabronero, J., Williams, G.B., Pereira, J.M.S., Pengas, G., Nestor, P.J., 2008. The impact of skull-stripping and radio-frequency bias correction on grey-matter segmentation for voxel-based morphometry. Neuroimage 39, 1654–1665.

Autio, J.A., Glasser, M.F., Ose, T., Donahue, C.J., Bastiani, M., Ohno, M., Kawabata, Y., Urushibata, Y., Murata, K., Nishigori, K., Yamaguchi, M., Hori, Y., Yoshida, A., Go, Y., Coalson, T.S., Jbabdi, S., Sotiropoulos, S.N., Kennedy, H., Smith, S., Van Essen, D.C., Hayashi, T., 2020a. Towards HCP-Style macaque connectomes: 24-Channel 3T multi-array coil, MRI sequences and preprocessing. Neuroimage 215, 116800.

Autio, J.A., Zhu, Q., Li, X., Glasser, M.F., Schwiedrzik, C.M., Fair, D.A., Zimmermann, J., Yacoub, E., Menon, R.S., Van Essen, D.C., Hayashi, T., Russ, B., Vanduffel, W., 2020b. Minimal Specifications for Non-Human Primate MRI: Challenges in Standardizing and Harmonizing Data Collection. arXiv: 2010.04325.

Avants, B.B., Epstein, C.L., Grossman, M., Gee, J.C., 2008. Symmetric diffeomorphic image registration with cross-correlation: evaluating automated labeling of elderly and neurodegenerative brain. Med. Image Anal. 12, 26–41.

Avants, B.B., Tustison, N., Song, G., 2009. Advanced normalization tools (ANTS). Insight J. 2, 1–35.

Benn, R.A., Mars, R.B., Xu, T., Rodríguez-Esparragoza, L., 2020. A Pig White Matter Atlas and Common Connectivity Space Provide a Roadmap for the Introduction of a New Animal Model in Translational Neuroscience. bioRxiv.

Bogart, S.L., Mangin, J.-F., Schapiro, S.J., Reamer, L., Bennett, A.J., Pierre, P.J., Hopkins, W.D., 2012. Cortical sulci asymmetries in chimpanzees and macaques: a new look at an old idea. Neuroimage 61, 533–541.

Buades, A., Coll, B., Morel, J.-M., 2011. Non-local means denoising. Image Processing On Line 1, 208–212.

Chen, W., Liu, B., Peng, S., Sun, J., Qiao, X., 2019. S3D-UNet: Separable 3D U-Net for Brain Tumor Segmentation. Brainlesion: Glioma, Multiple Sclerosis, Stroke and Traumatic Brain Injuries. https://doi.org/10.1007/978-3-030-11726-9_32

Cox, R.W., 1996. AFNI: software for analysis and visualization of functional magnetic resonance neuroimages. Comput. Biomed. Res. 29, 162–173.

Craddock, C., Sikka, S., Cheung, B., Khanuja, R., Ghosh, S.S., Yan, C., Li, Q., Lurie, D., Vogelstein, J., Burns, R., Others, 2013. Towards automated analysis of connectomes: The configurable pipeline for the analysis of connectomes (c-pac). Front. Neuroinform. 42.

Croxson, P.L., Forkel, S.J., Cerliani, L., Thiebaut de Schotten, M., 2018. Structural Variability Across the Primate Brain: A Cross-Species Comparison. Cereb. Cortex 28, 3829–3841.

Eskildsen, S.F., Coupé, P., Fonov, V., Manjón, J.V., Leung, K.K., Guizard, N., Wassef, S.N., Østergaard, L.R., Collins, D.L., Alzheimer’s Disease Neuroimaging Initiative, 2012. BEaST: brain extraction based on nonlocal segmentation technique. Neuroimage 59, 2362–2373.

Esteban, O., Markiewicz, C.J., Blair, R.W., Moodie, C.A., Isik, A.I., Erramuzpe, A., Kent, J.D., Goncalves, M., DuPre, E., Snyder, M., Oya, H., Ghosh, S.S., Wright, J., Durnez, J., Poldrack, R.A., Gorgolewski, K.J., 2019. fMRIPrep: a robust preprocessing pipeline for functional MRI. Nat. Methods 16, 111–116.

Fischl, B., 2012. FreeSurfer. Neuroimage 62, 774–781.

Fox, A.S., Oler, J.A., Birn, R.M., Shackman, A.J., Alexander, A.L., Kalin, N.H., 2018. Functional Connectivity within the Primate Extended Amygdala Is Heritable and Associated with Early-Life Anxious Temperament. J. Neurosci. 38, 7611–7621.

Fox, A.S., Oler, J.A., Shackman, A.J., Shelton, S.E., Raveendran, M., McKay, D.R., Converse, A.K., Alexander, A., Davidson, R.J., Blangero, J., Rogers, J., Kalin, N.H., 2015. Intergenerational neural mediators of early-life anxious temperament. Proc. Natl. Acad. Sci. U. S. A. 112, 9118–9122.

Ghafoorian, M., Mehrtash, A., Kapur, T., Karssemeijer, N., Marchiori, E., Pesteie, M., Guttmann, C.R.G., de Leeuw, F.-E., Tempany, C.M., van Ginneken, B., Fedorov, A., Abolmaesumi, P., Platel, B., Wells, W.M., 2017. Transfer Learning for Domain Adaptation in MRI: Application in Brain Lesion Segmentation. Medical Image Computing and Computer Assisted Intervention − MICCAI 2017. https://doi.org/10.1007/978-3-319-66179-7_59

Glasser, M.F., Sotiropoulos, S.N., Wilson, J.A., Coalson, T.S., Fischl, B., Andersson, J.L., Xu, J., Jbabdi, S., Webster, M., Polimeni, J.R., Van Essen, D.C., Jenkinson, M., WU-Minn HCP Consortium, 2013. The minimal preprocessing pipelines for the Human Connectome Project. Neuroimage 80, 105–124.

Henschel, L., Conjeti, S., Estrada, S., Diers, K., Fischl, B., Reuter, M., 2020. FastSurfer - A fast and accurate deep learning based neuroimaging pipeline. Neuroimage 219, 117012.

Hopkins, W.D., 2018. Motor and Communicative Correlates of the Inferior Frontal Gyrus (Broca’s Area) in Chimpanzees. Origins of Human Language: Continuities and Discontinuities with Nonhuman Primates 153.

Hopkins, W.D., Meguerditchian, A., Coulon, O., Bogart, S., Mangin, J.-F., Sherwood, C.C., Grabowski, M.W., Bennett, A.J., Pierre, P.J., Fears, S., Woods, R., Hof, P.R., Vauclair, J., 2014. Evolution of the central sulcus morphology in primates. Brain Behav. Evol. 84, 19–30.

Hwang, H., Rehman, H.Z.U., Lee, S., 2019. 3D U-Net for Skull Stripping in Brain MRI. Applied Sciences. https://doi.org/10.3390/app9030569

Isensee, F., Jaeger, P.F., Kohl, S.A.A., Petersen, J., Maier-Hein, K.H., 2021. nnU-Net: a self-configuring method for deep learning-based biomedical image segmentation. Nat. Methods 18, 203–211.

Jenkinson, M., Beckmann, C.F., Behrens, T.E.J., Woolrich, M.W., Smith, S.M., 2012. FSL. Neuroimage 62, 782–790.

Jung, B., Taylor, P.A., Seidlitz, J., Sponheim, C., Perkins, P., 2020. A comprehensive macaque fMRI pipeline and hierarchical atlas. NeuroImage. this issue.

Ketkar, N., 2017. Introduction to PyTorch. Deep Learning with Python. https://doi.org/10.1007/978-1-4842-2766-4_12

Kingma, D.P., Ba, J., 2014. Adam: A Method for Stochastic Optimization. arXiv [cs.LG].

Kleesiek, J., Urban, G., Hubert, A., Schwarz, D., Maier-Hein, K., Bendszus, M., Biller, A., 2016. Deep MRI brain extraction: A 3D convolutional neural network for skull stripping. Neuroimage 129, 460–469.

Lepage, C., Wagstyl, K., Jung, B., Seidlitz, J., Sponheim, C., Ungerleider, L., Wang, X., Evans, A.C., Messinger, A., 2021. CIVET-Macaque: An automated pipeline for MRI-based cortical surface generation and cortical thickness in macaques. Neuroimage 227, 117622. this issue.

Lohmeier, J., Kaneko, T., Hamm, B., Makowski, M.R., Okano, H., 2019. atlasBREX: Automated template-derived brain extraction in animal MRI. Sci. Rep. 9, 12219.

Lyksborg, M., Puonti, O., Agn, M., Larsen, R., 2015. An Ensemble of 2D Convolutional Neural Networks for Tumor Segmentation. Image Analysis. https://doi.org/10.1007/978-3-319-19665-7_17

McDonald, A.R., Muraskin, J., Van Dam, N.T., Froehlich, C., Puccio, B., Pellman, J., Bauer, C.C.C., Akeyson, A., Breland, M.M., Calhoun, V.D., Carter, S., Chang, T.P., Gessner, C., Gianonne, A., Giavasis, S., Glass, J., Homann, S., King, M., Kramer, M., Landis, D., Lieval, A., Lisinski, J., Mackay-Brandt, A., Miller, B., Panek, L., Reed, H., Santiago, C., Schoell, E., Sinnig, R., Sital, M., Taverna, E., Tobe, R., Trautman, K., Varghese, B., Walden, L., Wang, R., Waters, A.B., Wood, D.C., Castellanos, F.X., Leventhal, B., Colcombe, S.J., LaConte, S., Milham, M.P., Craddock, R.C., 2017. The real-time fMRI neurofeedback based stratification of Default Network Regulation Neuroimaging data repository. Neuroimage 146, 157–170.

Messinger, A., Sirmpilatze, N., Heuer, K., Loh, K.K., Mars, R.B., Sein, J., Xu, T., Glen, D., Jung, B., Seidlitz, J., Taylor, P., Toro, R., Garza-Villarreal, E.A., Sponheim, C., Wang, X., Benn, R.A., Cagna, B., Dadarwal, R., Evrard, H.C., Garcia-Saldivar, P., Giavasis, S., Hartig, R., Lepage, C., Liu, C., Majka, P., Merchant, H., Milham, M.P., Rosa, M.G.P., Tasserie, J., Uhrig, L., Margulies, D.S., Klink, P.C., 2021. A collaborative resource platform for non-human primate neuroimaging. Neuroimage 226, 117519. this issue.

Milham, M.P., Ai, L., Koo, B., Xu, T., Amiez, C., Balezeau, F., Baxter, M.G., Blezer, E.L.A., Brochier, T., Chen, A., Croxson, P.L., Damatac, C.G., Dehaene, S., Everling, S., Fair, D.A., Fleysher, L., Freiwald, W., Froudist-Walsh, S., Griffiths, T.D., Guedj, C., Hadj-Bouziane, F., Ben Hamed, S., Harel, N., Hiba, B., Jarraya, B., Jung, B., Kastner, S., Klink, P.C., Kwok, S.C., Laland, K.N., Leopold, D.A., Lindenfors, P., Mars, R.B., Menon, R.S., Messinger, A., Meunier, M., Mok, K., Morrison, J.H., Nacef, J., Nagy, J., Rios, M.O., Petkov, C.I., Pinsk, M., Poirier, C., Procyk, E., Rajimehr, R., Reader, S.M., Roelfsema, P.R., Rudko, D.A., Rushworth, M.F.S., Russ, B.E., Sallet, J., Schmid, M.C., Schwiedrzik, C.M., Seidlitz, J., Sein, J., Shmuel, A., Sullivan, E.L., Ungerleider, L., Thiele, A., Todorov, O.S., Tsao, D., Wang, Z., Wilson, C.R.E., Yacoub, E., Ye, F.Q., Zarco, W., Zhou, Y.-D., Margulies, D.S., Schroeder, C.E., 2018. An Open Resource for Non-human Primate Imaging. Neuron 100, 61–74.e2.

Oler, J.A., Fox, A.S., Shelton, S.E., Rogers, J., Dyer, T.D., Davidson, R.J., Shelledy, W., Oakes, T.R., Blangero, J., Kalin, N.H., 2010. Amygdalar and hippocampal substrates of anxious temperament differ in their heritability. Nature 466, 864–868.

Pontes-Filho, S., Dahl, A.G., Nichele, S., Gustavo Borges Moreno, 2019. A deep learning based tool for automatic brain extraction from functional magnetic resonance images in rodents. arXiv [eess.IV].

Puccio, B., Pooley, J.P., Pellman, J.S., Taverna, E.C., Craddock, R.C., 2016. The preprocessed connectomes project repository of manually corrected skull-stripped T1-weighted anatomical MRI data. Gigascience 5, 45.

Rehman, S., Ajmal, H., Farooq, U., Ain, Q.U., Riaz, F., Hassan, A., 2018. Convolutional neural network based image segmentation: a review. Pattern Recognition and Tracking XXIX. https://doi.org/10.1117/12.2304711

Ronneberger, O., Fischer, P., Brox, T., 2015. U-Net: Convolutional Networks for Biomedical Image Segmentation. Lecture Notes in Computer Science. https://doi.org/10.1007/978-3-319-24574-4_28

Roy, S., Butman, J.A., Reich, D.S., Calabresi, P.A., Pham, D.L., 2018. Multiple Sclerosis Lesion Segmentation from Brain MRI via Fully Convolutional Neural Networks. arXiv: 1803.09172.

Roy, S., Knutsen, A., Korotcov, A., Bosomtwi, A., Dardzinski, B., Butman, J.A., Pham, D.L., 2018. A deep learning framework for brain extraction in humans and animals with traumatic brain injury, in: 2018 IEEE 15th International Symposium on Biomedical Imaging (ISBI 2018). pp. 687–691.

Salehi, S.S.M., Hashemi, S.R., Velasco-Annis, C., Ouaalam, A., Estroff, J.A., Erdogmus, D., Warfield, S.K., Gholipour, A., 2018. Real-time automatic fetal brain extraction in fetal MRI by deep learning, in: 2018 IEEE 15th International Symposium on Biomedical Imaging (ISBI 2018). pp. 720–724.

Ségonne, F., Dale, A.M., Busa, E., Glessner, M., Salat, D., Hahn, H.K., Fischl, B., 2004. A hybrid approach to the skull stripping problem in MRI. Neuroimage 22, 1060–1075.

Seidlitz, J., Sponheim, C., Glen, D., Ye, F.Q., Saleem, K.S., Leopold, D.A., Ungerleider, L., Messinger, A., 2018. A population MRI brain template and analysis tools for the macaque. Neuroimage 170, 121–131.

Sørensen, T., 1948. A Method of Establishing Groups of Equal Amplitude in Plant Sociology Based on Similarity of Species Content and Its Application to Analyses of the Vegetation on Danish Commons.

Tasserie, J., Grigis, A., Uhrig, L., Dupont, M., Amadon, A., Jarraya, B., 2020. Pypreclin: An automatic pipeline for macaque functional MRI preprocessing. Neuroimage 207, 116353.

Tustison, N.J., Avants, B.B., Cook, P.A., Zheng, Y., Egan, A., Yushkevich, P.A., Gee, J.C., 2010. N4ITK: improved N3 bias correction. IEEE Trans. Med. Imaging 29, 1310–1320.

Tustison, N.J., Cook, P.A., Holbrook, A.J., Johnson, H.J., Muschelli, J., Devanyi, G.A., Duda, J.T., Das, S.R., Cullen, N.C., Gillen, D.L., Others, 2020. ANTsX: A dynamic ecosystem for quantitative biological and medical imaging. medRxiv.

Xu, T., Sturgeon, D., Ramirez, J.S.B., Froudist-Walsh, S., Margulies, D.S., Schroeder, C.E., Fair, D.A., Milham, M.P., 2019. Interindividual Variability of Functional Connectivity in Awake and Anesthetized Rhesus Macaque Monkeys. Biol Psychiatry Cogn Neurosci Neuroimaging 4, 543–553.

Xu, T., Yang, Z., Jiang, L., Xing, X.-X., Zuo, X.-N., 2015. A Connectome Computation System for discovery science of brain. Sci Bull. Fac. Agric. Kyushu Univ. 60, 86–95.

Yan, C.-G., Wang, X.-D., Zuo, X.-N., Zang, Y.-F., 2016. DPABI: Data Processing & Analysis for (Resting-State) Brain Imaging. Neuroinformatics 14, 339–351.

Yogananda, C.G.B., Wagner, B.C., Murugesan, G.K., Madhuranthakam, A., Maldjian, J.A., 2019. A Deep Learning Pipeline for Automatic Skull Stripping and Brain Segmentation, in: 2019 IEEE 16th International Symposium on Biomedical Imaging (ISBI 2019). pp. 727–731.

Yosinski, J., Clune, J., Bengio, Y., Lipson, H., 2014. How transferable are features in deep neural networks?, in: Ghahramani, Z., Welling, M., Cortes, C., Lawrence, N., Weinberger, K.Q. (Eds.), Advances in Neural Information Processing Systems. Curran Associates, Inc., pp. 3320–3328.

Yushkevich, P.A., Piven, J., Hazlett, H.C., Smith, R.G., Ho, S., Gee, J.C., Gerig, G., 2006. User-guided 3D active contour segmentation of anatomical structures: significantly improved efficiency and reliability. Neuroimage 31, 1116–1128.

Zhao, G., Liu, F., Oler, J.A., Meyerand, M.E., Kalin, N.H., Birn, R.M., 2018. Bayesian convolutional neural network based MRI brain extraction on nonhuman primates. Neuroimage 175, 32–44.

